# Brain lipidomics identifies mitochondrial redox dysfunction and metabolic trade-offs associated with Parkinson’s disease-like pathology induced by Nanoplastics exposure

**DOI:** 10.1101/2025.11.09.687439

**Authors:** Priya Rathor, Ashutosh K. Tiwari, Rajendra P Patel, Sheelendra Pratap Singh, Ratnasekhar CH

**Author notes:** Corresponding author: E-mail addresses.

## Abstract

Growing nanoplastics exposure raises concern for neurotoxicity, particularly given recent evidence of plastic accumulation within human brain tissue-a highly lipid enriched organ, yet effects on brain lipid metabolism remains poorly understood. Here, we employed high-resolution untargeted lipidomics to map brain lipid perturbations in *Drosophila melanogaster* chronically exposed to environmentally relevant levels of polystyrene nanoplastics (NPs). Polystyrene NPs accumulated in fly brains and induced dose-dependent remodeling of mitochondrial membrane lipids, notably cardiolipins and phosphatidylethanolamines, accompanied by increased diacylglycerols/triacylglycerols and monounsaturated fatty acids and by lipid droplet expansion. Guided by these lipidomic signatures, targeted biochemical assays demonstrated depolarized mitochondrial membrane potential, elevated mitochondrial reactive-oxygen species, inhibition of respiratory-chain complexes I and IV, and a shift in NAD(H) and NADP(H) redox couples toward a reduced state and increasing lipid peroxidation. This redox imbalance was accompanied by decreased tyrosine-hydroxylase expression, dopamine depletion, and impaired locomotor behavior, hallmarks of PD-like neurodegeneration. Dopaminergic neurochemistry was impaired (tyrosine hydroxylase and dopamine decreased), with concomitant reduction of GABA, and locomotor and circadian deficits emerged. Remarkbly, co-treatment with the antioxidant N-acetylcysteine (NAC) restored mitochondrial membrane potential, reduced mitochondrial ROS and lipid peroxidation, normalized neutral lipid and MUFA accumulation, and rescued neurotransmitter levels and behavior. Stable-isotope tracing confirmed disrupted TCA cycle flux after NP exposure that was rescued by NAC. Collectively, these findings reveal lipidomic remodeling as a critical link between environmental nanoplastic exposure and PD-like pathology, highlighting mitochondrial redox–lipid interactions as early determinants and support redox-directed interventions to mitigate risk.

## 1. Introduction

Over recent decades, the relentless accumulation of plastic waste in terrestrial and aquatic environments has emerged as a critical global concern, giving rise to a new class of pollutants microplastics (MPs) and nanoplastics (NPs) [1]. Despite their minuscule dimensions, NPs pose disproportionate biological risks due to their small size (<200nm), chemical stability, high surface reactivity, and capacity to interact with living systems at the cellular and molecular levels [2,3]. With global plastic production projected to surpass 1.2 billion tonnes by 2060 and recycling rates remaining alarmingly low, the environmental and biological burdens of micro- and nanoplastics are intensifying at an unprecedented scale [4]. These particles have now permeated virtually every environmental compartment such as food, drinking water, and even the air facilitating multiple human exposure routes including ingestion, inhalation, and dermal contact. This raises urgent concerns for both ecological balance and human health, particularly about their potential health impacts [3,5,6].

The neurological impacts of NPs represent a particularly concerning area of toxicity. Recent analytical breakthroughs have confirmed that NPs not only enter peripheral tissues (lungs, placenta, kidneys) but also accumulate in the brain, with concentrations markedly higher among individuals suffering from neurodegenerative conditions such as dementia and Alzheimer’s disease [7,8]. This discovery has intensified concerns about the potential neurotoxic effects of NPs, especially as both laboratory and epidemiological studies now link MNPs exposure to cognitive dysfunction and increased risk for neurological disorders [7–9]. While early research focused on the systemic and peripheral toxicity of NPs, a growing body of animal and cellular studies reveal that NPs can cross the blood–brain barrier (BBB) via multiple routes, breach neural tissues, and disrupt the homeostasis of neurons and glia. The olfactory bulb has been identified as a plausible entry point, enabling MPs to bypass the BBB and potentially access deeper brain structures. Once inside the brain, these particles can induce oxidative stress, neuroinflammation, and apoptosis, all critical pathological mechanisms underlying neurodegeneration [10]. Studies in rodent models demonstrate that chronic NPs exposure leads to cognitive deficits [11]. An expanding body of preclinical evidence implicates NPs in the disruption of neural development and brain physiology [12]. Recent study has shown that anionic NPs can form high-affinity complexes with the amphipathic and non-amyloid component domains of α-synuclein, identifying a potential mechanistic link between NPs exposure and PD–related synucleinopathies [13].

Moreover, behavioral abnormalities have been observed across diverse aquatic species, including nematodes, crustaceans, and fish, following NPs exposure. Structural and functional impairments of nerve fibers, along with deficits in locomotor activity and enzymatic functions, have also been documented. High-resolution human postmortem studies further highlight the urgent need to understand the consequences of cerebral plastic accumulation, as dementia cases not only have greater plastic burden but may also be exposed to compounding processes such as barrier breakdown and reduced clearance. Taken together, these findings underscore the neurotoxic potential of NPs and highlight the urgent need for studies to assess their long-term effects on brain health.

The brain is a lipid-rich organ, with nearly 50–60% of its dry weight composed of complex lipids that are essential for membrane architecture, synaptic transmission, and signaling. Even subtle perturbations in brain lipid metabolism can disrupt neuronal communication, mitochondrial function, and neuroinflammatory balance, thereby predisposing to neurodegenerative processes [14]. In the current neurotoxicological paradigm, such lipid metabolic disturbances can act as early molecular events preceding overt pathology, particularly under environmental stressors like NPs exposure. Lipidomics the comprehensive analysis of lipidomes using high-resolution accurate mass spectrometry (HRAMS) enables offering a sensitive readout of the brain’s metabolic state. In this context, high-throughput MS-based lipidomics can uncover alterations in lipid metabolites that are critical for neuronal membrane integrity, neurotransmitter release, and oxidative stress regulation. Moreover, cumulative shifts in lipid composition and distribution in response to external stressors like NPs can provide valuable insights into the physiological status, and lipids themselves can act as critical regulators of cellular function and phenotype [15–20]. Despite the brain’s vulnerability to environmental stressors, NPs-induced perturbations of endogenous lipid species in this organ remain not explored. There is a major knowledge gap regarding the impact of NPs exposure on brain lipid metabolism which are increasingly recognized as central to the pathogenesis of neurodegenerative diseases such as Parkinson’s. Given the central role of lipids in maintaining neuronal function and the potential for lipid dysregulation to contribute to neurotoxicity, in-depth lipidomic investigation is warranted to elucidate these molecular alterations [21].

The present study explored the NPs-induced toxic effects on brain lipid metabolism and associated pathways in *Drosophila melanogaster*, a well-established experimental model that shares ∼75% of human disease-related genes, exhibits high conservation with the human brain, making it a powerful system for modelling neurotoxic effects. Owing to these genetic and functional similarities, *Drosophila* has been extensively used in dissecting brain-specific metabolic processes. Coupling this model with high-resolution lipidomics enables the characterization of conserved lipid pathways disrupted by environmental stressors, NPs thereby providing mechanistic insights with direct translational relevance to human neurotoxicity. *Drosophila* was exposed to environmentally relevant concentrations of NPs (0.05 mg/mL). In addition, a low-concentration (0.01 mg/mL) exposure group was designed. Untargeted lipidomics was performed to assess the impact of NPs exposure on brain lipid metabolism. Global lipidomic analysis reveals specific alterations in mitochondrial lipids phosphatidylethanolamine (PE) and cardiolipin (CL). Disruption in mitochondrial lipids precedes accumulation of lipid droplets with increased diglyceride (DG) and triglyceride (TG) lipids. To confirm the mitochondrial vulnerability, mitochondrial membrane potential (MMP) and mitochondrial reactive oxygen species (mtROS) were determined. We further examine whether these changes are associated with key neurochemical alterations such as deficits in dopamine metabolism and whether antioxidant intervention can mitigate both molecular and behavioral disruptions. By integrating lipidomic, bioenergetic, and stable isotope–resolved metabolomic analyses, we aimed to establish a link between NPs-induced mitochondrial dysfunction, impaired central carbon metabolism, and NAC-mediated metabolic rescue.

## 2. Materials and methods

### 2.1 Chemicals and reagents

Acetonitrile (Optima LC-MS grade, Sigma-Aldrich), methanol, ethanol, isopropanol, and chloroform (LC-MS grade, Sigma-Aldrich,St. Louis, MO, USA) were used in this study. Ultra-pure water (LC-MS grade) was obtained from Sigma-Aldrich. 4-(2-Hydroxyethyl) piperazine-1-ethanesulfonic acid (HEPES), sodium dodecyl sulfate (SDS), Hydrogen peroxide (H□O□) was purchased from Himedia. Dichloro-dihydro-fluorescein diacetate (DCFH-DA) was obtained from Cayman Chemicals, while thiobarbituric acid (TBA) and tetraethoxypropane (TEP) were purchased from Tokyo Chemical Industry (Japan). JC-1 dye and MitoSOX dye were obtained from Thermo Fisher Scientific. The Complex I enzyme activity assay kit was purchased from Abcam. NAD/NADH and NADP/NADPH quantification kits were procured from Sigma-Aldrich. Prodan and BODIPY dyes were purchased from Tokyo Chemical Industry (Japan). Acetic acid was obtained from TCI, sodium phosphate from Sigma-Aldrich, nitroblue tetrazolium (NBT) from RealGene, and phenazine methosulfate from TCI. The Bradford protein assay kit was purchased from Thermo Fisher Scientific. Phosphate-buffered saline (PBS) was obtained from Himedia. Dimethyl sulfoxide (DMSO) was purchased from Sigma-Aldrich, and Triton X-100 was obtained from Himedia.

### 2.2 Nanoplastics and *Drosophila* culture

Nonfluorescent and fluorescent polystyrene-NPs suspensions were purchased from, nanoshell and alpha nanotech respectively. NPs were characterized by using transmission electron microscopy (TEM), Dynamic light scattering (DLS), Raman and Fourier transform-infrared (FT-IR) spectroscopy. Wild-type *Drosophila* (*Oregon R+)* was cultured under standardized laboratory conditions, maintained at 25°C with 60% relative humidity and a 12-hour light/dark photoperiod to ensure consistency across all experimental groups. To minimize environmental and handling variability, all experiments were conducted under these controlled conditions. The nutritional composition per litre included 8 g agar, 15 g yeast, 80 g cornmeal, 20 g sucrose, 10 g glucose, and 1 g methylparaben as a mold inhibitor. Unless stated otherwise, only male flies were used in experiments to reduce variability due to sex-specific metabolic differences [22 23]. PS-NPs beads size 100 nm was evenly mixed into the agar-based fly diet before solidification. Flies were randomly distributed into three groups including control, fed a sucrose-agar medium and treatment groups. The NPs exposure concentrations (0.01 mg/ml and 0.05 mg/ml) were chosen based on environmentally relevant doses reported in prior studies [24,25]. Four days old adult flies were transferred to NPs-supplemented diets and maintained on these media for seven consecutive days to assess dose-dependent brain lipidomic, physiological and biochemical responses. For validation experiments, *Oregon R*□ flies were treated with N-acetylcysteine (NAC) at a previous reported concentration of 1 mg/mL, following the same protocol as used for NPs treatment [26].

### 2.3 NPs brain internalization

Fluorescent polystyrene-NPs were obtained from Alpha Nanotech (Green fluorescent polymer microspheres, λ_ex: 488 nm, λ_em: 518 nm, particle size100 nm, 10 mg/mL suspension, USA). To ensure homogeneous distribution, the required concentration of NPs was incorporated into agar-based diet prior to solidification. The prepared media were wrapped in aluminium foil and stored in the dark to prevent photobleaching. Experimental *Drosophila* were transferred to NPs-supplemented diets containing 0.01 mg/ml and 0.05 mg/ml and maintained under these conditions for seven days to assess dose-dependent effects. For brain imaging, fly heads were dissected and fixed in 4% paraformaldehyde at room temperature for 24 h. Following fixation, the head cuticle was carefully removed, and brains were mounted on glass slides using 70% glycerol. A coverslip was applied and sealed with transparent nail polish to prevent tissue rupture. Brains were then imaged using confocal microscopy (Zeiss LSM 880) under green fluorescence settings (λ_ex: 488 nm, λ_em: 518 nm). Fluorescence intensity was quantified using ImageJ software [27].

### 2.4 Histopathological evaluation of *Drosophila* brain

Histological analysis of *Drosophila* brains was performed using standard hematoxylin and eosin (H & E) staining. Adult fly heads were dissected in 1× PBS and fixed overnight at 4 °C in 4% paraformaldehyde. Fixed samples were dehydrated through a graded ethanol series (30%, 50%, 70%, 90%, and 100%) and cleared with xylene before embedding in paraffin wax. Paraffin blocks were sectioned at 5 µm thickness using a rotary microtome, and sections were mounted onto wax-coated glass slides. The sections were deparaffinized in xylene, rehydrated through descending ethanol concentrations, and stained with hematoxylin for 5 minutes. Following are rinsed in water, differentiation was performed using acid alcohol, and sections were blued in alkaline water. Subsequently, counterstaining was carried out with eosin for 2 minutes, followed by dehydration, clearing, and mounting in 80% glycerol. Stained brain sections were examined under a bright-field light microscope (Leica), and images were captured for comparative evaluation of brain morphology between control and NPs treated groups [28].

### 2.5 Scanning Electron Microscopy (SEM) analysis of brain samples

Adult fly heads were dissected in 1× PBS and immediately fixed in 1 mL of freshly prepared 4% paraformaldehyde (PFA) in PBS. Samples were incubated overnight at 4 °C to ensure optimal tissue preservation. Following fixation, the PFA solution was discarded, and tissues were washed three times with 0.1% PBST (PBS containing 0.1% Tween-20) at room temperature for 10 minutes each to remove residual fixative. Brains were then carefully dissected from the fixed heads in 1× PBS and subjected to an additional three washes in 0.1% PBST under the same conditions to eliminate surface contaminants. Dehydration was performed through a graded ethanol series (30%, 50%, 70%, 90%, and 100%), with each step lasting 10 minutes and repeated twice to ensure complete dehydration. After the final wash, samples were air-dried thoroughly. The dried brains were mounted onto aluminum SEM stubs using conductive adhesive and sputter-coated with a thin layer of conductive metal (gold or platinum) to improve imaging quality. SEM imaging was performed under high-vacuum conditions using standard operating parameters to visualize NPs-induced ultrastructural changes in the fly brain tissue [29].

### 2.6 Membrane polarity assessment using Prodan dye

Fly brains were dissected in ice-cold PBS and incubated with 5 μM Prodan (6-propionyl-2-(dimethylamino) naphthalene) (diluted from DMSO in PBS) for 30 minutes at room temperature in the dark. Following staining, samples were washed three times with PBS to remove unbound dye and mounted in PBS. Imaging was performed using fluorescence or confocal microscopy with excitation at 360 nm and dual emission acquisition at 440 nm and 520 nm. The emission intensity ratio reflected membrane polarity changes, with a blue shift indicating increased hydrophobicity.

### 2.7 Determination of lipid droplets in *Drosophila* brain tissue

Brains from adult flies were dissected in ice-cold 1XPBS and rinsed three times with 1× PBS. The tissues were then incubated for 10 minutes in 0.1% Triton X-100 in PBS to facilitate dye penetration. Subsequently, the brains were stained with 2 µM BODIPY dye for 30 minutes at room temperature under dark conditions. Following staining, samples were washed three times with PBS to remove excess dye and mounted on glass slides using antifade Vectashield mounting medium without DAPI. Imaging was recorded with excitation at 405 nm and emission at 520 nm [30].

### 2.8 Extraction of lipids from *Drosophila* brains

After seven days of NPs exposure, *Drosophila* samples were rapidly quenched in liquid nitrogen, and brain tissues were dissected. Lipid extraction was carried out using a modified protocol from Xue Li Guan et al. [31].The dissected brain tissues were homogenized in 100 µL of ice-cold 1× PBS using a handheld motorized pestle. Briefly, 500 µL of chloroform: methanol (1:2, v/v) was added to the homogenate, vortexed for 1 min, and incubated in a Thermomixer (Eppendorf type C) at 1,000 rpm for 2 h at 4 °C. Subsequently, 200 µL of chloroform and 200 µL of water was added, and the mixture was centrifuged at 9,000 rpm for 2 min at 4 °C (Eppendorf 5424R). The lower organic phase was carefully collected and dried using a vacuum concentrator, and stored at -80^0^C until further use.

### 2.9 Comprehensive lipidome profiling of *Drosophila* brain samples using HRAMS

The dried brain tissue lipid extracts were reconstituted in 200 µL of precooled acetonitrile: methanol (1:1, v/v), and lipid separation was performed using reversed-phase liquid chromatography with slight modifications to a previously described method [32]. Lipidomic profiling was carried out on a Vanquish UHPLC system coupled to an Orbitrap Exploris 240 mass spectrometer (Thermo Scientific). LC separation was followed by using established protocols [33]. Briefly, a Zorbax Eclipse Plus C18 column (2.1 × 150 mm, 1.8 µm particle size) was employed with a binary gradient of solvent A (water: acetonitrile, 40:60, v/v) and solvent B (isopropanol: acetonitrile, 90:10, v/v), both supplemented with 10 mM ammonium acetate and 0.1% formic acid. A 27-min elution program was applied as follows: 0–3 min, 30% B; 5 min, 43% B; 5.1 min, 55% B; 10 min, 65% B; 12.5–13.5 min, 85% B; 18–23 min, 100% B; followed by re-equilibration at 30% B until 27 min. The LC parameters including flow rate 300 µL/min, column temperature 40 °C, injection volume 2 µL, and autosampler temperature 4 °C was used. Mass spectrometry was conducted using a HESI II probe in both positive and negative ion modes. The MS settings were full scan range 100–1500 m/z; resolution 120,000; spray voltage 3.8 kV (positive mode) and 3.5 kV (negative mode); probe temperature 330 °C; sheath gas 35 (arbitrary units); and auxiliary gas 10 (arbitrary units). Data were acquired using Xcalibur 4.5 software (Thermo Scientific). Data-dependent MS/MS analysis employed higher-energy collisional dissociation (HCD) with normalized collision energies of 20, 35, and 60. The pooled QC samples was run between the sample to check the sample acquisition reproducibility of instrument.

### 2.10 Untargeted lipidomics data processing

The data analysis was conducted as in our previous studies. For global lipidomics data extraction, the RAW data acquired from Xcalibur 4 were processed for peak detection and alignment using MSDIAL version 4.9.2. All steps in the MSDIAL lipidome data processing were carried out with previously described parameters, with minor modifications [34]. This afforded a list of data (mass-to-charge ratio, retention time, and peak intensity) in CSV format. The metabolic features of quality control samples with a standard deviation greater than 30% relative and disturbed signals in the blanks were excluded to eliminate interference. The MS/MS data were further processed for metabolite identification, which involved matching the retention time, accurate precursor mass, isotope pattern, and MS/MS spectra of the lipid metabolites to those of metabolite libraries. The metabolite libraries used were LIPID MAPS, LipidHome, LipidBlast, and *Drosophila* lipidome reference libraries. The molecular weight tolerance was set at ±5 ppm with respect to the theoretical values for each metabolite. The qualified data were analyzed for Multivariate statistical analysis was performed using principal component analysis (PCA) and partial least squares discriminant analysis (PLS-DA) to assess group differences. The PLS-DA model was built with R software packages. A heatmap was generated based on the abundance of differential lipids using R software. Correlation analysis was used to calculate the correlation coefficient (r) between different lipid groups. The following criteria were applied for univariate analysis to identify differential lipids: the Variable Importance for the Projection (VIP) in the first two components of the PLS-DA model was set to ≥ 1. Fold Changes (FCs) were categorized as upregulated if FC ≥ 1.5 and downregulated if FC ≤ 1.5.

### 2.11 Mitochondrial membrane potential (ΔΨm) analysis of fly brains

MMP was assessed using JC-1 dye as described previously [35]. Adult fly brains from NPs-treated, NAC-treated, and control groups were dissected in ice-cold PBS under a stereomicroscope. Dissected tissues were incubated with 5 µM JC-1 (prepared in DMSO) for 30 min at room temperature in the dark to minimize photobleaching. After incubation, brains were washed three times with 1XPBS (2 min each) to remove unbound dye and reduce background fluorescence. Stained samples were mounted in PBS on glass slides and imaged using a Zeiss LSM 880 confocal laser scanning microscope. JC-1 aggregates (red fluorescence, indicating polarized mitochondria) and JC-1 monomers (green fluorescence, indicating depolarized mitochondria) were detected using appropriate excitation/emission settings (red channel: Ex ∼540–560 nm, Em ∼570–590 nm). Quantitative analysis of mitochondrial polarization was performed by measuring the red fluorescence intensity using ImageJ software, with values normalized to control samples. All experiments were performed in three independent biological replicates to ensure reproducibility.

### 2.12 Mitochondrial ROS determination

Mitochondria were isolated from ∼100 adult fly heads homogenized in 1 mL of ice-cold isolation buffer (0.25 M Tris-sucrose, pH 7.4) containing 220 mM mannitol, 68 mM sucrose, 10 mM KCl, 10 mM HEPES, and 0.1% BSA. The homogenate was centrifuged twice at 300 × g to remove debris, followed by centrifugation at 3,000 × g for 10 min at 4 °C to pellet mitochondria. The mitochondrial pellet was washed and resuspended in isolation buffer without BSA. Mitochondrial ROS production was quantified using 2′,7′-dichlorodihydrofluorescein diacetate (DCFH-DA), final concentration 5 µM; stock prepared in DMSO. Fluorescence was recorded at 37 °C in a plate reader with excitation/emission settings of 488/520 nm, measured every minute for 10 min. Results were expressed as arbitrary fluorescence units (AFU) [36].

For mitochondrial superoxide (O^2-^) detection, adult fly brains were dissected using a stereomicroscope in ice-cold 1XPBS (at pH 7.4) following brief anaesthesia using ice. Brains were incubated with freshly prepared MitoSOX™ Red (Thermo Fisher Scientific, Cat. No. M36008; 5 µM in PBS) for 30 min at room temperature in the dark with gentle agitation. After incubation, tissues were washed three times with PBST (2 min each) to remove excess dye. Fluorescence signals were visualized using a TRITC filter set (Ex: ∼510 nm, Em: ∼580 nm). Image acquisition parameters were kept constant across all groups, and mean fluorescence intensity was quantified using ImageJ software [37].

### 2.13 Total ROS estimation in *Drosophila* brain samples

ROS levels in *Drosophila* brain samples were quantified following a previously described protocol with minor modifications [38].. Briefly, adult fly brain were dissected using ice-cold PBS and incubated with 10 μM 2′,7′-dichlorodihydrofluorescein diacetate (DCFH-DA) for 30 min at room temperature in the dark to minimize photobleaching. After incubation, tissues were fixed with 4% PFA to preserve fluorescence and structural integrity. The samples were then mounted on glass slides using 70% glycerol as the mounting medium. Fluorescence signals were visualized using a confocal microscope with identical acquisition settings maintained across all experimental groups. Quantification of ROS levels was performed by measuring mean fluorescence intensity using ImageJ software. All experiments were conducted in triplicate with independent biological replicates to ensure reproducibility and statistical reliability [38].

### 2.14 Mitochondrial Complex I activity assay

Mitochondrial complex I activity was measured in brain tissue samples using a commercial kit (Abcam, ab109721) with slight modifications [39]. Fresh brain tissues were rinsed with ice-cold PBS, homogenized in 500 µL chilled PBS on ice, and clarified lysates were obtained. Protein concentration was determined and normalized to 5 mg/mL in PBS. To solubilize mitochondrial membranes, one-tenth volume of supplied detergent was added and incubated for 30 min on ice, followed by centrifugation (16,000 × g, 20 min, 4 °C). Supernatants were loaded into pre-coated assay plate wells, washed, and activity was measured kinetically at 450 nm for 30 min at room temperature (at 1 min intervals). Results were expressed as the rate of absorbance change (mOD/min).

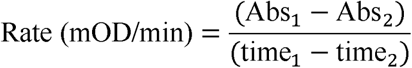

### 2.15 Mitochondrial cytochrome C activity assay

Cytochrome C oxidase activity was measured using the cytochrome C oxidase assay kit (Abcam, ab239711) following the manufacturer’s instructions. Freshly prepared brain tissue lysates were homogenized in cold PBS and kept on ice throughout processing, including centrifugation. Reduced cytochrome C was prepared by reconstituting the reagent with assay buffer, adding DTT, and incubated for 15 min at room temperature. A 1:6 dilution of reduced cytochrome C was made using pre-warmed assay buffer, and 120 µL was prepared per reaction. For the assay, 5–10 µL of brain tissue lysate and dilution buffer (blank) was added to each well of a 96-well plate. Then 120 µL of diluted reduced cytochrome C was added to initiate the reaction. Absorbance at 550 nm was recorded at 30-second intervals for 30 minutes using a kinetic program. Enzyme activity was calculated from the linear portion of the curve as ΔOD per minute (mOD/min):

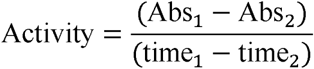

All measurements were performed in triplicate, and results were expressed relative to control groups [40].

### 2.16 Estimation of NAD**□**/NADH

The NAD□/NADH ratio in *Drosophila* brain tissue was determined using the NAD□/NADH Colorimetric Assay Kit (Sigma-Aldrich, Cat. No. MAK468),with the following the manufacturer’s protocol [41]. Briefly, ∼100 adult fly brain per group were homogenized with extraction buffer and centrifuged to obtain the supernatant. Heat treatment (at 60 °C, 30 min) was applied to differentiate NAD□ from NADH. Absorbance was measured at 565 nm using a microplate reader over a 15-min kinetic reading, and concentrations were calculated from a standard curve. The NAD□/NADH ratio was then determined for each experimental group.

### 2.17 Estimation of NADP**□**/NADPH

The NADP□/NADPH ratio in fly brain tissue was determined using the NADP□/NADPH Quantification Kit (Sigma-Aldrich, Cat. No. MAK479) with the manufacturer’s instructions [42]. Briefly, ∼100 adult fly brains were homogenized in the provided extraction buffer. NADPH levels were measured after heat treatment (60 °C, 30 min), while total NADP□ + NADPH levels were obtained from untreated samples. NADP□ concentration was calculated by subtracting NADPH from the total pool. Absorbance was recorded at 565 nm using a microplate reader, and concentrations were determined from a standard curve. Data were normalized to protein content, and NADP□/NADPH ratios were calculated for each group.

### 2.18 Lipid peroxidation assay

Lipid peroxidation was quantified by measuring malondialdehyde (MDA), a stable end-product of oxidative degradation of lipids, using a modified thiobarbituric acid reactive substances (TBARS) assay [43]. Tetraethoxypropane was used as an external reference for calibration. In brief, 25 μL of 10% fly brain homogenate was mixed with 150 μL of 10% SDS, 1.0 mL of 0.8% thiobarbituric acid, and 1.0 mL of 20% acetic acid (at pH 3.5). The mixture was incubated in a boiling water bath at 100 °C for 60 min to allow color development, cooled to room temperature, and extracted with 3.0 mL of n-butanol. Absorbance was measured at 535 nm using n-butanol as the blank. The assay was performed in five independent biological replicates with duplicate technical measurements. MDA levels were determined from the standard calibration curve and normalized to protein concentration, expressed as nmol per mg of protein [44].

### 2.19 Catalase Activity Assay

Catalase activity was determined following the method described previously [45], with slight modifications. The assay is based on the ability of catalase to decompose hydrogen peroxide (H□O□). Reagents prepared included dichromate–acetic acid, 0.02 M H□O□, and 0.01 M sodium phosphate buffer. For the assay, 1 mL of phosphate buffer was transferred into a 15 mL falcon tube, followed by deionized water. Subsequently, 25 μL of a 10% tissue homogenate was added. The mixture was gently vortexed, after which 0.5 mL of 0.02 M H□O□ was introduced and mixed again. The reaction was terminated by adding 2.0 mL of dichromate–acetic acid, and the tubes were incubated in a boiling water bath (100 °C) for 15 minutes. After cooling, absorbance was recorded at 570 nm using a spectrophotometer. Each experiment was performed independently five times, with samples run in triplicate. Data were normalized to protein concentration and expressed as micromoles of H□O□ decomposed per milligram of protein (µmol H□O□/mg protein).

### 2.20 Superoxide dismutase (SOD) assay

SOD activity in the brain of NPs-exposed *Drosophila* was determined using Beauchamp and Fridovich method [46]. The assay mixture contained freshly prepared sodium pyrophosphate buffer, nitroblue tetrazolium (NBT), NADH, and phenazine methosulfate (PMS). Tissue homogenates were prepared from precisely 20 brains per sample for both protein quantification and enzyme activity measurement. The enzymatic reaction was initiated by adding 25□µL of the homogenate to the reaction mixture, and the reduction of NBT was monitored spectrophotometrically at 560□nm over a 20-minute period. SOD activity was defined as the amount of protein required to inhibit the NBT reduction rate by 50% under the assay conditions. At least four independent biological replicates were assessed per experimental group.

### 2.21 Estimation of protein

Protein content was estimated using the Bradford assay [47]. Briefly, 20 brain samples were homogenized in buffer (0.1M sodium phosphate, pH 7.4) and centrifuged at 8,000 × g for 5 min at 4 °C. A 10 µL aliquot of the resulting supernatant was mixed with 490 µL of Bradford reagent. For the blank, an equal volume of homogenization buffer was used in place of the sample or standard. The reaction mixtures were incubated at room temperature in the dark for 30 min, and absorbance was measured at 595 nm. Protein concentration (mg/µL) was determined from the standard calibration curve.

### 2.22 Estimation of GABA and dopamine following NPs exposure

Precisely, 100 fly brains were collected and homogenized with precooled 80% methanol (v/v) using a pestle-mortar followed by sonication for 15 min under ice-cold conditions to ensure efficient metabolite extraction. The brain tissue homogenates were subsequently vortexed for an additional 15 min to enhance solubilization of intracellular metabolites. The samples were then centrifuged at 14,000 rpm for 15 min at 4 °C, and the supernatants were carefully collected. The extracts were dried using a vacuum concentrator and reconstituted in LC-MS grade 80% methanol (v/v). The reconstituted samples were subjected to HRAMS analysis for the relative quantification of γ-aminobutyric acid (GABA) and dopamine with selective ion monitoring (SIM) in positive polarity mode.

### 2.23 Immunofluorescence analysis of dopaminergic neurons

To visualize dopaminergic neurons, immunofluorescence staining was carried out on isolated brain tissues of control and NPs-exposed flies, following a protocol was adapted from earlier described protocol [48].Adult fly brains were dissected in ice-cold PBS and fixed in 4% paraformaldehyde, followed by several washes with PBS (0.01 M PBS containing 0.1% Triton X-100). The samples were then incubated overnight at 4 °C in blocking buffer (PBST supplemented with 5% heat-inactivated fetal bovine serum) to minimize nonspecific binding. After blocking, brains were incubated with rabbit polyclonal anti-tyrosine hydroxylase (TH) antibody (1:200; Abcam) overnight at 4 °C. The following day, excess primary antibody was removed with PBST washes, and tissues were incubated for 2 h at room temperature (24 ± 2 °C) with a Cy3-conjugated goat anti-rabbit secondary antibody (1:250; Abcam). After secondary labeling, brains were rinsed with PBS, mounted in Vectashield antifade medium (Vector Laboratories, CA, USA), and imaged using a Leica confocal microscope (Nussloch, Germany). The number of TH-positive neurons within major dopaminergic clusters was quantified by examining confocal Z-series projections.

### 2.24 Tyrosine hydroxylase (TH) activity assay

TH activity was assessed using a previously described protocol adapted from Vermeer et al. [49] with modifications as described by Musachio et al. [50]. Briefly, flies were cryo-anesthetized and decapitated. For each biological replicate (n), twenty fly heads were homogenized in 250 μL of Tris-HCl buffer (0.05 M, pH 7.2). The homogenates were centrifuged at 13,000 × g for 5 min at 4 °C, and 100 μL of the supernatant was mixed with an equal volume of reaction buffer containing 100 mM HEPES, 100 μM tyrosine, and 200 μM sodium periodate. The enzymatic reaction was monitored spectrophotometrically at 475 nm for 30 min at 25 °C. TH activity was expressed as a percentage relative to the control group. All assays were performed in five independent experiments (n = 5).

### 2.25 Uniform 13C-Glucose tracing and measured the glycolytic and TCA cycle intermediate metabolites alteration

Following seven days of NPs exposure, adult *Drosophila* were subjected to a 6-hour starvation period to deplete internal energy reserves. Subsequently, flies from both control and treated groups were administered a uniform ¹³C□-glucose (U-¹³C□, >99% atom, Cambridge Isotope Laboratories) diet for 48 hours to trace metabolic flux through central carbon metabolism according to previous reported study [51]. Post-treatment, flies were harvested and snap-frozen in liquid nitrogen, followed by metabolite extraction using chilled 80% methanol (v/v). Each sample underwent triple extraction, and pooled supernatants were lyophilized and reconstituted in LC-MS-grade MeOH prior to analysis. Metabolomic profiling was performed using a Thermo Scientific Vanquish UHPLC system coupled with an Orbitrap Exploris 240 mass spectrometer (Waltham, MA, USA). To maximize metabolite coverage, dual injections of 10 µL were analyzed in negative ion mode, and an additional 2 µL injection was analyzed in positive ion mode. Chromatographic separation was achieved under hydrophilic interaction liquid chromatography (HILIC) conditions using a Waters XBridge BEH Amide column (150 × 2.1 mm, 2.5 µm particle size). The mobile phase was delivered at a flow rate of 0.3 mL/min, with the autosampler maintained at 4°C and the column oven at 40°C. LC parameters were consistent with those used in our targeted metabolomics workflow to ensure cross-sample comparability. Mass spectrometric data were acquired in full scan mode (m/z 70–1,000) using electrospray ionization (ESI). Data acquisition and peak extraction were performed using Thermo Scientific software and processed further for isotopologue analysis. Metabolite identification and isotopic labeling patterns were annotated by matching features to an in-house spectral library of ∼600 reference standards, supplemented by the HMDB, METLIN, and other commercial databases. Annotation criteria included exact mass (±5 ppm), retention time, isotopic distribution, and MS/MS fragmentation, with a minimum absolute intensity threshold of 1,000 counts. Glycolysis and TCA cycle metabolites were the primary focus for assessing ¹³C incorporation patterns and metabolic pathway activity.

### 2.26 Locomotor activity monitoring

Locomotor behavior was assessed using the *Drosophila* Activity Monitoring (DAM2) system (TriKinetics, USA). Flies aged 2–3 days were maintained on a standard cornmeal diet at 25 °C before the assay. Individual flies were gently introduced into glass tubes (65 mm × 5 mm) containing food at one end, prepared from 5% sucrose and 1.2% agar. The tubes were placed in the DAM2 system under a controlled 12 h light/dark cycle at 25 °C. Following a 24 h entrainment period, baseline activity was recorded for two consecutive days. On day three, immediately after lights-on, flies were transferred to fresh food tubes containing either 0.01 mg/mL and 0.05 mg/mL NPs, or unsupplemented medium (control). Locomotor activity was then continuously monitored for an additional seven days. Data were binned into 30 min intervals for the entire recording period. To evaluate NPs-induced effects, locomotor activity on day seven was compared with baseline activity levels. Data processing was performed using DAMFileScan113 software, and statistical analyses were conducted with GraphPad Prism 8.4.2 (TriKinetics; https://www.trikinetics.com/).

### 2.27 Climbing assay

After exposure to NPs, locomotor activity of treated and control flies was evaluated using a negative geotaxis climbing test [52]. For each group, 20 flies were placed in a vertical plastic column (18 cm in height × 2 cm in diameter) and allowed to acclimate for 1 min. The flies were then gently tapped to the bottom, and their climbing ability was recorded over a 30 s period. Flies that crossed the 15 cm mark and those that remained below it were counted separately. Each assay consisted of five consecutive trials with 1min intervals between trials. A performance index (PI) was calculated using the formula: PI = ½[(Ntot + Ntop − Nbot)/Ntot], where Ntop denotes the number of flies above the 15 cm mark, Nbot represents flies below the mark, and Ntot is the total number of flies tested. Data were obtained from three independent experiments and are reported as mean ± standard deviation (SD).

### 2.28 Statistical analysis

All descriptive data were analyzed using R (V 4.4.2) and GraphPad Prism (version 8.1) software. ANOVA followed by turkey’s multiple comparison test was used to evaluate the statistical differences. The results from data are presented as mean ± standard error of mean (SEM), with n =6 biological replicates. Statistical significance was represented by *p* < 0.05.

## 3. Results and discussion

Understanding the potential neurotoxic effects of NPs is crucial, as humans are continuously exposed to these particles through food, water, and air. Once internalized, NPs may reach the brain, an organ in which lipids constitute more than half of its dry weight and play indispensable roles in neuronal function, signaling, and structural integrity. In this context, we employed *Drosophila* as a well-established model organism to evaluate the effects of PS-NPs, 100 nm on brain lipid metabolism, providing a tractable framework to understand how environmental NPs exposure could contribute to neurological risks.

### 3.1 Brain internalization of NPs, and its effects on morphology, membrane integrity and accumulation of lipid droplets

We first characterized of PS-NPs and then used for downstream biological experiments in *Drosophila*. High-magnification TEM revealed that the PS-NPs exhibited predominantly spherical morphology (Fig. S1A). Size distribution analysis by intensity using DLS demonstrated a sharp and narrow peak centered around ∼100 nm, confirming the mono-dispersity of the NPs (Fig. S1B). Raman spectroscopy and FT-IR analysis revealed prominent consistent with the functional groups typically found in polystyrene. These results confirm the presence of polymeric bonds and verify the structural composition of the NPs (Fig.S1C & D). Additionally, we performed the confocal imaging of fluorescent labelled NPs (Fig.S1E). Internalization assays with and without fluorescent labelling were conducted to investigate whether NPs entered the brain. *Drosophila* flies were treated with environmentally relevant concentrations of 0.01 mg/mL and 0.05 mg/mL. An experimental plan for PS-NPs exposure is represented in Fig 1A. The fluorescent NPs were found to accumulate in the brain in a dose-dependent manner. NPs uptake was evident in the higher NPs concentration groups (0.05 mg/mL) (Fig. 1B). This observation is consistent with earlier reports showing that NPs can cross biological barriers and accumulate in neural tissues, including the BBB in zebrafish and mice [53,54].

**Figure 1:**
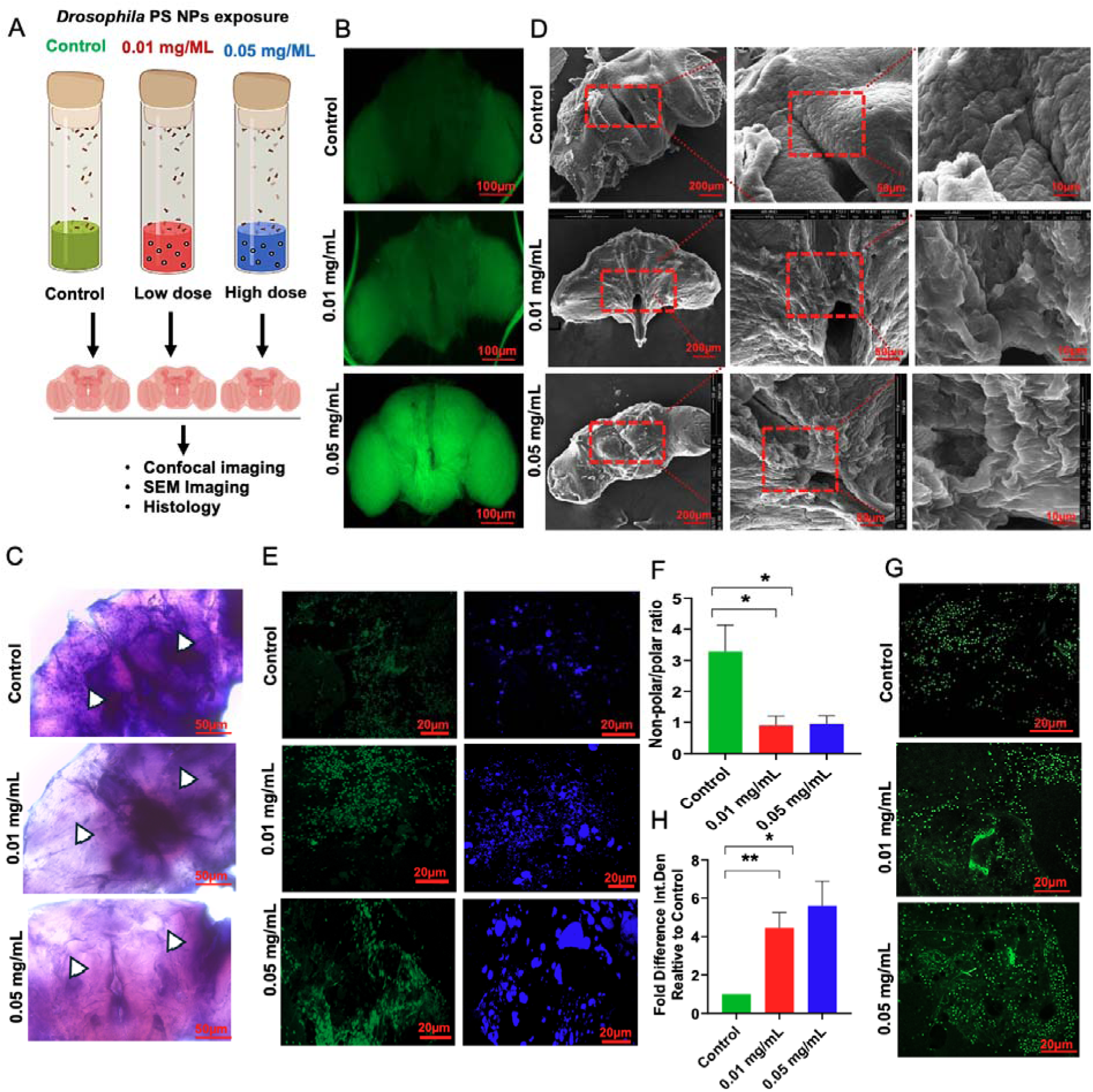
Neuro-accumulation and neuronal disruption in *Drosophila melanogaster* upon NPs exposure. **(A).** Experimental plan for PS-NPs treatment of flies**, (B)** Confocal images of fluorescent-labeled NPs accumulate in brains **(C)** H&E-stained brain sections exhibit increased vacuolization (white arrowheads) in NPs-exposed brains, especially at 0.05 mg/mL. **(D)** SEM images of *Drosophila* brain at increasing magnifications (200 µm, 50 µm, and 10 µm) reveal surface deformation damage in treated groups. **(E-F)** Confocal images of Prodan-stained *Drosophila* brains. Green fluorescence marks non-polar membrane regions (intact polarity), while blue fluorescence indicates increased membrane polarity (stress or damage). NPs-treated groups show enhanced blue fluorescence, indicating membrane depolarization and structural compromise. **(G-H)** Confocal images of BODIPY stained *Drosophila* brains, which selectively label neutral lipid droplets. Treated groups display increased green fluorescence, denoting enhanced lipid droplet accumulation often associated with oxidative or metabolic stress. Scale bars = 20□μm. Values are expressed as Mean ± Standard Error of Mean (n = 6). Significant differences from the control are indicated *p* <* 0.05, *p**<*0.005, *p****<*0.0001, student T-Test.

H&E staining was performed to assess the histoarchitecture of *Drosophila* brains following NPs exposure. In the control group the brain tissue appeared well-organized, with distinct cellular regions and compact neuropil. There were no signs of tissue disintegration or vacuolization. In contrast, brains from flies exposed to 0.01 mg/mL NPs exhibited moderate structural disruption, with loosening of brain architecture and the appearance of faint vacuolated regions (indicated by white arrowheads). This indicates early signs of neurodegenerative changes. Exposure to 0.05 mg/mL NPs (Fig.1C), resulted in severe neurodegeneration, evident by prominent vacuole formation, diffused cellular outlines, and tissue disintegration throughout the brain region. Similar histopathological features have been noted in rodent models exposed to PS-NPs [55]. The density and clarity of neuropil were markedly reduced, suggesting progressive loss of neuronal integrity. These results were consistent with previous results that significant pathological changes were observed in the brain tissue, including neuronal crumpling, intensified staining, and the formation of small vacuoles in polystyrene MPs-induced zebrafish (Danio rerio) [56].

To further evaluate whether NPs exposure induces structural alterations in neural tissue, we performed SEM imaging on dissected *Drosophila* brains (Fig. 1D). In control flies, the brain surface appeared smooth and well-organized. In contrast, flies exposed to 0.01 mg/mL NPs exhibited early surface irregularities, while higher concentrations (0.05 mg/mL) led to pronounced architectural disorganization, including surface collapse, roughening, and loss of structural integrity. These progressive alterations reflect tissue-level disruption and suggest that NPs accumulation compromises membrane stability. These morphological changes may compromise membrane stability and prompted us to assess membrane integrity more directly. Given this evidence, we next investigated membrane integrity directly using Prodan dye staining. Consistent with the SEM observations, Prodan staining revealed a concentration-dependent disruption of membrane integrity in NPs-exposed brains. At low NPs concentrations, fluorescence remained largely uniform, whereas higher concentrations elicited markedly enhanced and irregular staining patterns (Fig. 1E & F), reflecting compromised membrane organization.

Given that membrane disruption and remodeling are closely linked to lipid homeostasis, we next investigated whether NPs exposure promotes lipid droplet accumulation as part of the adaptive cellular response. Consistent with this, BODIPY staining revealed a dose-dependent increase in lipid droplet accumulation in NPs-exposed fly brains, with semi-quantitative analysis showing a significant elevation in staining intensity following exposure (*p* < 0.05; Fig.1G & H). Supporting our findings, previous studies have reported lipid droplet accumulation in the intestines of mice exposed to NPs, suggesting that NPs exposure induces lipid droplet accumulation [17]. Lipid droplets act as reservoirs for neutral lipids and typically expand under conditions of metabolic stress [57,58]. Notably, the brain is composed of more than 50% lipids, making it particularly vulnerable to NP-induced lipid perturbations. Considering this, NPs exposure is likely to disrupt the lipidome of the brain. Therefore, we further investigated whether NPs exposure damages brain lipid structures and sought to elucidate the underlying mechanisms of NPs-induced lipid disruption.

### 3.2 Comprehensive brain lipidome profiling identify endogenous lipid biomarkers in responsed to NPs exposure

Next, we asked how the *Drosophila.* brain lipidome is perturbed upon chronic exposure of NPs at different concentrations. To address this, we performed global lipidomic profiling of brain tissue using LC-HRAMS. Total brain lipid extracts were analyzed in DDA mode (Fig. 2A). The tight clustering of pooled QC samples demonstrated excellent reproducibility of the acquired lipidomic data. Lipids were identified based on exact mass measurements with sub-p.p.m. accuracy, elemental composition constraints, and further validated by MS/MS fragmentation of the corresponding precursors.

**Figure 2:**
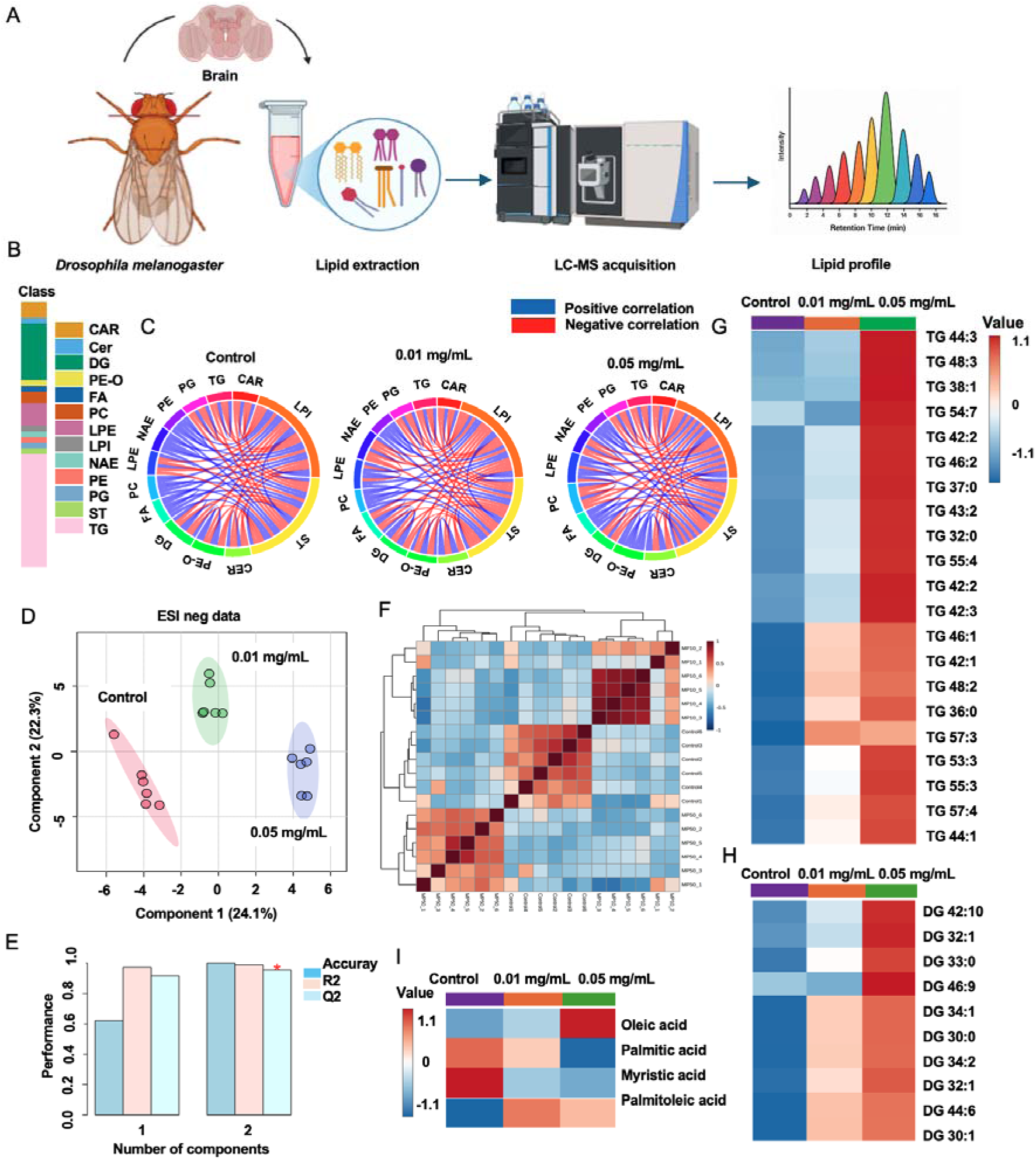
NPs induced brain lipidome alteration (**A).** Schematic represents the brain lipidome study design (**B).** Investigated some lipid subclasses in brain tissue under NPs treated condition. (**C).** Chord diagrams showing the intra-tissue correlation of lipids classes within brain tissue with 0.01 and 0.05 mg/mL NPs**. (D)** PLS-DA components accounted for 46.4% of the variance (**E).** PLS-DA demonstrated strong performance with cumulative values of R² = 0.98597, Q² = 0.95145, and Accuracy = 1 **(F).** Pearson’s correlation **(G-I)** Heatmap plot for identified TG and DG lipids and fatty acids subclasses in brain tissue.

Across all datasets, we systematically identified 357 lipids spanning diverse categories including fatty acyls (FA), glycerolipids (GL), glycerophospholipids (GP), sphingolipids (SL), and sterol lipids (ST) in brain tissue extracts (Fig S2). Scatter plots depict the ionization-dependent coverage of lipid classes in both ESI (+) and ESI (−) modes (Fig. S3). GP species predominated in the negative mode (Fig. S3A), whereas abundant GL and FA signals were detected in the positive mode (Fig. S3B). Sub-classification into 13 functional lipid classes, such as DG, TG, PE, LPI, PG, CL, PEO, PC, LPE, ST, FA, NAE, Cer, and CAR, revealed distinct patterns of NPs-induced alterations, summarized in Fig. 2B. Circos analysis of 13 lipid classes revealed that control brains displayed predominantly positive correlations, reflecting balanced coordination between membrane phospholipids (PE, PC, PG) and neutral lipids (DG, TG). At 0.01 mg/mL NPs exposure, correlation density increased with stronger positive interactions (e.g., DG–PC, TG–PE) but also the emergence of negative associations involving lysophospholipids (LPE, LPI), indicating early adaptive remodeling of lipid metabolism and droplet expansion under stress, summarized in Fig. 2C. At 0.05 mg/mL, negative correlations became more widespread, particularly between carnitines and glycerophospholipids and between lysophospholipids and neutral lipids, pointing to mitochondrial lipid transport defects and membrane turnover imbalance.

Multivariate analysis was employed to distinguish the lipid profiles across sample classes. Partial least squares discriminant analysis (PLS-DA) was employed to identify the lipid species contributing to the separation between control and NPs-exposed *Drosophila* under both ESI (−) and ESI (+) ionization modes (Fig 2D). The corresponding score plots displayed clear and distinct clustering of individual samples within their respective groups across both ionization modes. To remove any overfitting, internal cross-validation was performed. In the ESI (−) mode, the PLS-DA model exhibited excellent performance, with cumulative values of R² = 0.98597, Q² = 0.95145, and classification accuracy of 1.0 (Fig. 2E). To further validate the robustness of the models, permutation testing was conducted using 500 permutations. This analysis confirmed the high predictive power and reliability of the optimized PLS-DA models in discriminating NPs-induced samples from controls, with statistical significance established at P < 0.01 (Fig.S3C). In addition, Pearson correlation analysis of the control and NPs-treated brain extracts revealed strong positive correlations within each group, underscoring the reproducibility and consistency of the datasets (Fig. 2F). Significant lipids were identified as key discriminatory features distinguishing control from NPs-treated groups based on the variable importance in projection (VIP) scores (VIP > 1) derived from the PLS-DA models, and P < 0.05 (Table S1).

Global lipidomic analysis revealed significant perturbations in neutral lipid metabolism, most prominently characterized by robust elevation in DG and TG lipid pools. Both DG and TG are critical intermediates in the neutral lipid pathway, where DG functions as a biosynthetic precursor and TG represents the primary constituent of lipid droplets (Fig. 2G & H). The progressive accumulation of DG and TG indicates activation of lipid droplet biogenesis as a compensatory cellular response to NPs-induced stress. Consistent with this interpretation, a dose-dependent increase in lipid droplet formation was visualized in NPs-exposed brains (Fig. 1 G & H), mirroring the elevated TG levels and confirming lipid storage adaptation under NPs challenge. Because DG serves as a precursor in fatty acid metabolism, facilitating TG synthesis and lipid droplet expansion, we next examined fatty acid composition in NPs-exposed brains. This was accompanied by a significant enrichment of monounsaturated fatty acids (MUFAs), particularly oleic acid and palmitoleic acid, compared with controls (Fig. 2I). MUFAs are known to be preferentially incorporated into triacylglycerols and stored in lipid droplets as a protective mechanism against excess ROS and lipid peroxidation. The concurrent increase in MUFA content and lipid droplet accumulation therefore suggests an adaptive cellular response aimed at buffering NPs-induced oxidative stress and preserving membrane integrity. However, persistent and excessive lipid droplet biogenesis may disrupt cellular lipid homeostasis, impair mitochondrial–ER communication, and contribute to neurodegenerative progression. Importantly, similar alterations have been described in PD models driven by α-synuclein toxicity, where elevated DG, TG, and MUFA levels were identified as part of a maladaptive lipid remodeling program that promotes lipid droplet accumulation and neuronal stress [59]. In these models, stress-induced MUFA accumulation serves a redox-protective function but, when chronically activated, contributes to α-synuclein aggregation, vesicular trafficking defects, and neurodegeneration. Our findings parallel these observations, suggesting that NPs exposure elicits PD-like lipidomic signatures, linking altered neutral lipid and MUFA metabolism with hallmarks of synucleinopathy. Together, these results highlight that NPs exposure triggers pronounced perturbations in DG and TG metabolism, leading to enhanced lipid droplet biogenesis and remodeling of fatty acid composition in the *Drosophila* brain. These lipidomic alterations likely represent an adaptive yet maladaptive metabolic response, wherein lipid remodeling contributes to disturbed membrane architecture, altered organelle function, and exacerbated cellular stress, thereby providing mechanistic insight into NPs-induced neurotoxicity.

In addition to TG and DG lipid alterations, we found perturbations in levels of mitochondrial-associated marker lipids, particularly CL and PE, in NPs-exposed brain tissue suggested potential compromise in mitochondrial membrane structure and function (Fig S4A & B). Previous studies on liver cells have suggested that mitochondria is sensitive to damage from NPs, as mitochondria are crucial to maintain cellular function. Therefore, we further explored whether NPs exposure damaged mitochondria function.

### 3.3 NPs exposure damages brain mitochondrial metabolism

Mitochondria are central regulators of neuronal energy homeostasis, and their membranes are enriched in CL and PE, two critical lipids that maintain respiratory chain integrity (Fig.3A & B) and modulate oxidative stress responses [60,61]. Given that our lipidomics analysis revealed perturbations in these classes, we hypothesized that NPs exposure may disrupt mitochondrial function through membrane damage (Fig 3C & D). Because CL and PE are tightly linked to mitochondrial membrane architecture, alterations in their abundance may reflect destabilization of the inner mitochondrial membrane, leading to impaired MMP [62]. To test this, we assessed MMP in *Drosophila* brains using JC-1, a potentiometric dye widely used to monitor mitochondrial health [63]. In healthy cells, high drives JC-1 aggregation, yielding dominant red fluorescence, whereas in depolarized mitochondria the dye remains in its monomeric form, producing increased green fluorescence [63]. A dose-dependent shift from red to green fluorescence was observed in NPs-exposed brains (0.01 and 0.05 mg/mL), indicating progressive mitochondrial depolarization and loss of bioenergetic integrity (Fig 3E & F).

**Figure 3.**
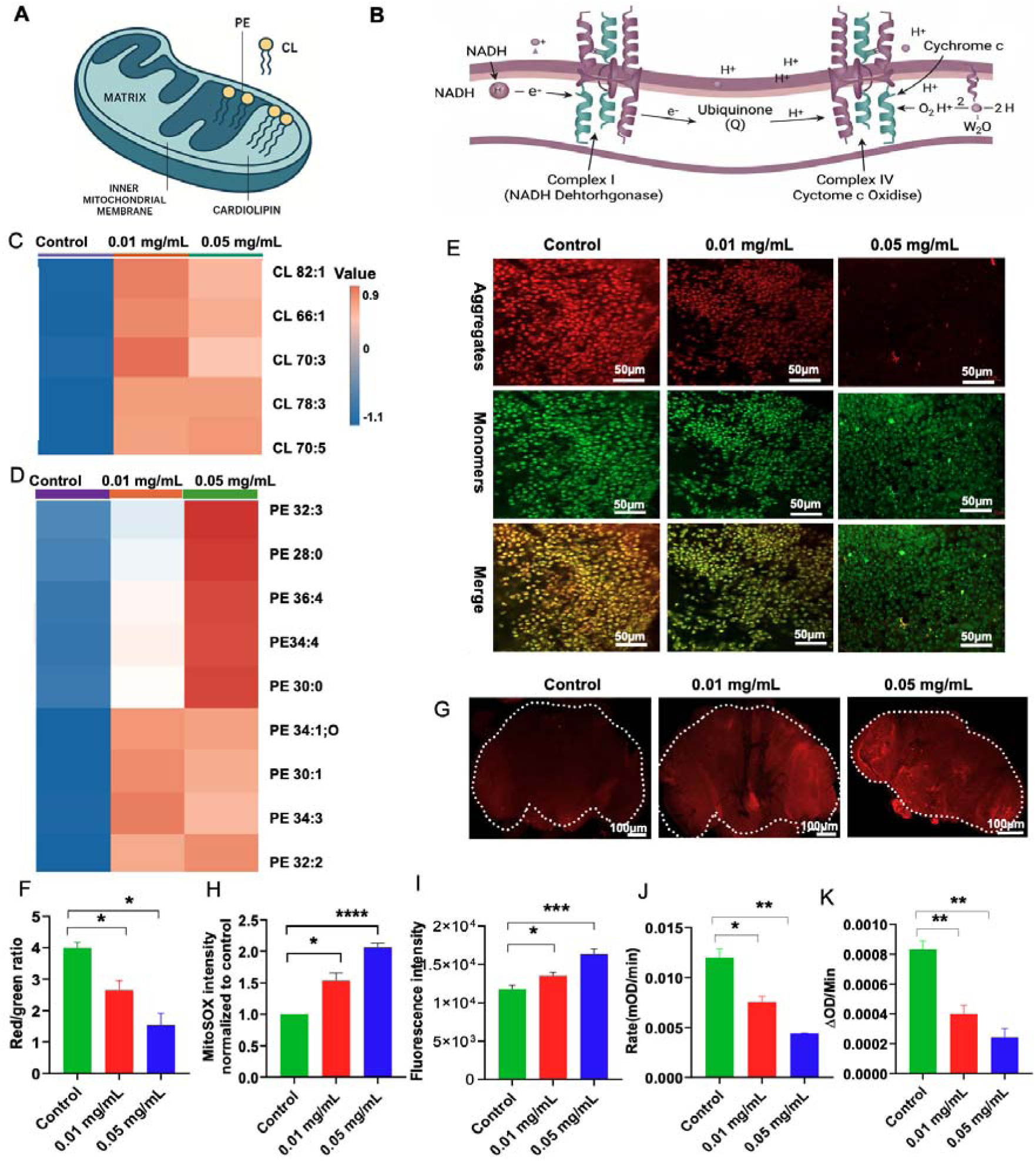
NPs exposure alters mitochondrial membrane lipids, impairs mitochondrial function, and elevates oxidative stress in the *Drosophila* brain. **(A)** Schematic of mitochondrial lipids. **(B)** Mitochondrial electron chain transport **(C&D)** Heatmap of significantly altered cardiolipin **(CL)** and phosphatidylethanolamine **(PE)** species in the brain following 0.01 and 0.05 mg/mL NPs exposure. Elevated levels of CL 82:1, CL 66:1, CL 70:3, CL 78:3, and CL 70:5 suggest dysregulation of mitochondrial membrane composition and PE 32:3, PE 28:0, PE 36:4, PE 34:4, PE 30:0, PE 34:1; O, and others, indicating perturbation of cellular membrane lipid profiles. Results are visualized in a color scale where blue indicate a decrease in values and red an increase. Heatmap was normalized from − 1.1 to 0.9 **(E&F)** JC-1 staining of *Drosophila* brain mitochondria reveals a dose-dependent loss of mitochondrial membrane potential under NPs exposure. Red (J-aggregates) indicates healthy polarized mitochondria, while green (monomers) reflects depolarized mitochondria. A decrease in red/green fluorescence ratio at 0.05 mg/mL indicates significant mitochondrial dysfunction. Scale bar = 50 μm. JC-1 red/green fluorescence ratio showing reduced mitochondrial membrane potential. **(G&H)** Representative confocal images of MitoSOX-stained *Drosophila* brains display enhanced red fluorescence in the NPs-treated groups, indicating elevated mitochondrial superoxide accumulation. Brains exposed to 0.05 mg/mL NPs exhibit the most intense and widespread MitoSOX signal. Scale bar = 100 µm**. (I)** Mitochondrial ROS levels are elevated, suggesting increased oxidative burden **(J)** Complex I activity significantly declines in a dose-dependent manner. **(K)**. Cytochrome C oxidase activity significantly declines in a dose-dependent manner. Values are expressed as Mean ± Standard Error of Mean (n = 5). Significant differences from the control are indicated p* < 0.05, p**<0.005, p****<0.0001 Student t-test.

Because excessive mtROS are a well-established driver of mitochondrial depolarization and membrane damage [64,65], we next investigated whether NPs exposure enhances oxidative stress. Using MitoSOX Red, a fluorogenic dye specific for mitochondrial superoxide (O2-), we found a marked increase in red fluorescence intensity in NPs-exposed groups, with the strongest signal in the 0.05 mg/mL group. Quantitative analysis confirmed a dose-dependent elevation of mtROS levels, supporting the conclusion that NPs exposure induces oxidative stress mediated mitochondrial dysfunction (Fig 3G & H). To further validate the role of oxidative stress in NPs-induced mitochondrial dysfunction, we quantified mtROS using DCFH-DA fluorescence in isolated brain mitochondria. Consistent with our MitoSOX results, DCFH-DA analysis revealed a dose-dependent increase in ROS levels, with significant changes detectable even at 0.01 mg/mL NPs exposure (Fig 3I). Fluorescence imaging of *Drosophila* brains confirmed this trend, showing progressive ROS accumulation after NPs. Quantitative analysis demonstrated significantly elevated fluorescence intensity in both the 0.01 mg/mL and 0.05 mg/mL groups compared with controls (p < 0.001; Fig. 3H). Together with MitoSOX staining, these findings provide robust evidence that NPs ingestion triggers excessive mtROS generation, reinforcing oxidative stress as a central mechanism of NPs-induced mitochondrial damage. These findings are consistent with previous reports demonstrating that NPs-treated L02 liver cells exhibit elevated levels of mtROS [63]. These results suggest that NPs exposure contributed to the generation of a oxidative stress environment in *Drosophila* brain.

The altered MMP and elevated mtROS observed in our study are likely linked to impaired electron transport chain (ETC) activity, as disruptions of ETC complexes are known drivers of mitochondrial dysfunction [65]. To investigate this, we assessed the activities of complexes I and IV, which play critical roles in oxidative phosphorylation and respiratory efficiency. NPs exposure caused a dose-dependent decline in the activities of both complexes, with the most pronounced inhibition observed at 0.05 mg/mL in *Drosophila* brain samples (Fig 3J & K). Consistent with impaired mitochondrial respiration, NPs exposure caused a dose-dependent decline in NAD□ and NADP□ levels with a concomitant increase in NADH and NADPH, indicating a shift toward a more reduced redox state (Fig S5A-D). This redox imbalance correlated with enhanced MDA and total ROS accumulation and reduced SOD and catalase activities, confirming oxidative stress (Fig S6A-D). These results indicate that NPs exert a strong inhibitory effect on mitochondrial respiration, thereby reducing electron flux and compromising ATP production. Importantly, inhibition of complexes I and IV also provides a mechanistic basis for the observed increase in mtROS and collapse of membrane potential, as electron leakage from dysfunctional complexes promotes redox imbalance, and antioxidant depletion [64]. Similar associations between NPs exposure and ETC perturbations have been reported in human lung and liver cells, supporting the generalizability of our findings across tissues and species [66,63].

### 3.4 NPs-induced alterations in dopamine, and Tyrosine hydroxylase expression

Because mitochondrial dysfunction in brain is a hallmark of PD pathology and directly impacts neurotransmission, particularly in dopaminergic systems, we next examined whether NPs exposure alters neurotransmitter homeostasis in *Drosophila*. Quantification of dopamine levels revealed a significant reduction in NPs-exposed flies compared with controls, with the strongest decline at 0.05 mg/mL (Fig. 4A). This observation is consistent with PD-related neurodegeneration, where mitochondrial impairment and oxidative stress lead to dopaminergic vulnerability and dopamine depletion [67,68]. To further evaluate dopaminergic regulation, we measured TH, the rate-limiting enzyme in dopamine biosynthesis. NPs exposure caused a dose-dependent downregulation of TH expression (Fig.4B & C), paralleling the decline in dopamine levels. Reduced TH expression is a well-recognized PD hallmark and reflects impaired dopamine synthesis under conditions of mitochondrial dysfunction and oxidative damage [69,70]. To explore the mechanistic basis of dopamine loss, we measured tyrosine hydroxylase (TH), the rate-limiting enzyme in dopamine biosynthesis. NPs exposure resulted in concentration dependent downregulation of TH expression (Fig. 4D), paralleling the reduction in dopamine levels. Downregulation of TH has similarly been reported under conditions of mitochondrial impairment and oxidative stress, reinforcing the link between NPs-induced mitochondrial dysfunction and disrupted dopamine synthesis [69,70]. In parallel, we assessed GABA, the principal inhibitory neurotransmitter in the brain. GABA levels were also significantly reduced in NPs-exposed flies brain tissue (Fig. 4E), suggesting that NPs-induced mitochondrial dysfunction not only compromises excitatory dopaminergic signaling but also disrupts inhibitory neurotransmission. Altered GABAergic transmission is increasingly recognized as another PD-feature, contributing to perturbations of the excitatory–inhibitory balance are known to underlie neuronal vulnerability in models of neurotoxicity and PD [71].

**Figure 4:**
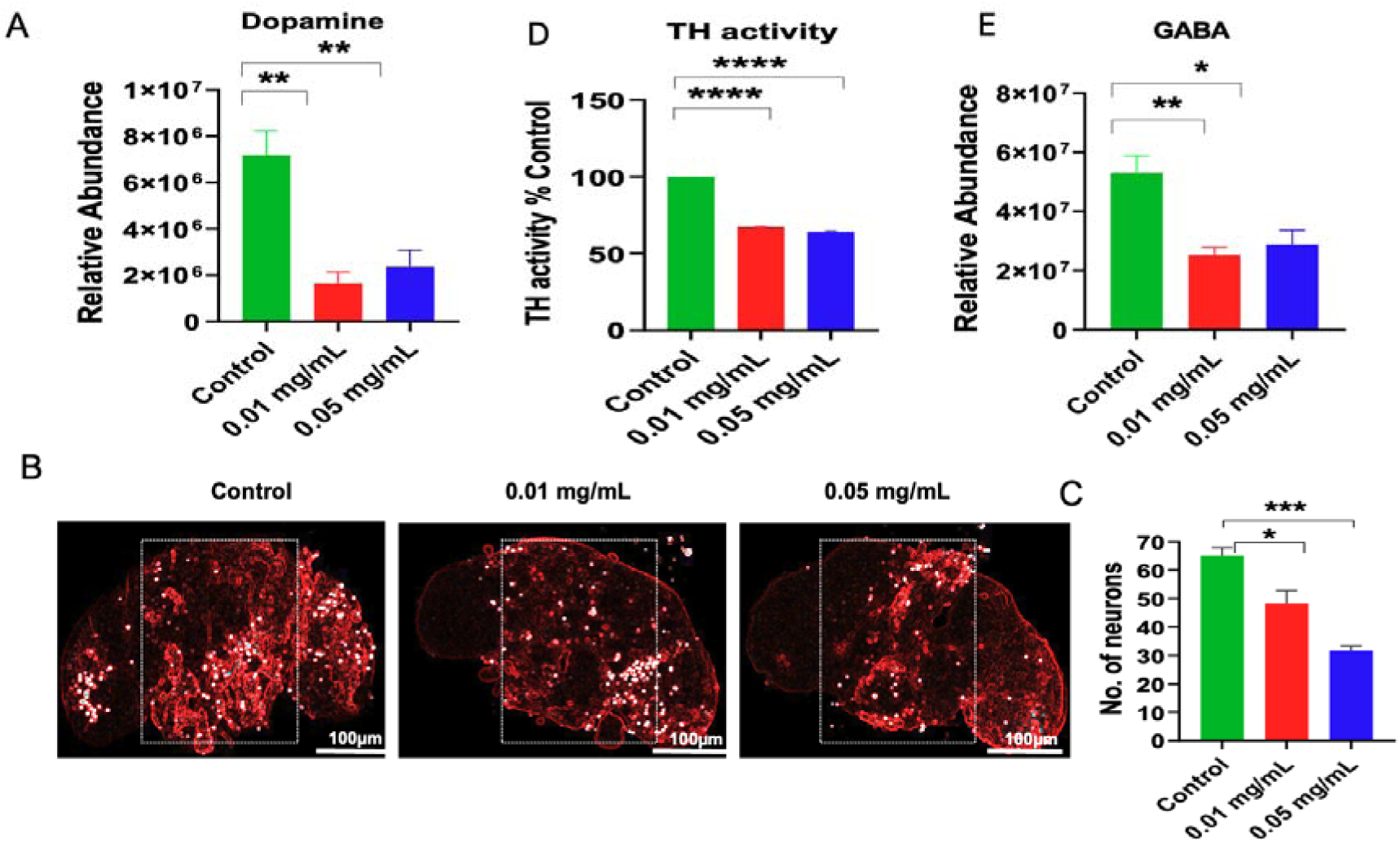
NPs-induced significant alteration of brain tissue specific neurotransmitters (**A)** Decreased the Dopamine level. **(B-C)** number of TH-positive neuron loss after the NPs exposure. **(D).** Measured Tyrosine hydrolase activity **(E)** Decreased the GABA level Values are expressed as Mean ± Standard Error of Mean (n = 5). Significant differences from the control are indicated p* < 0.05, p**<0.005, p****<0.0001 Student t-test.

Together, these findings demonstrate that NPs exposure disrupts mitochondrial bioenergetics and redox balance, which in turn leads to reduced dopamine and GABA levels, accompanied by downregulation of TH expression. These neurochemical alterations provide a mechanistic explanation for the behavioral impairments observed in NPs-exposed flies and are consistent with previous reports linking mitochondrial dysfunction to neurotransmitter dysregulation in neurodegenerative contexts.

### 3.5 Effect of chronic NPs exposure on *Drosophila* behavior and survival

The observed reductions in dopamine and GABA levels, together with downregulation of tyrosine hydroxylase expression, strongly suggest that NPs exposure disrupts excitatory and inhibitory neurotransmission. Such neurochemical imbalances are known to impair motor coordination, circadian regulation, and reduced neuronal resilience. To determine whether these neurochemical alterations translated into measurable phenotypic outcomes, we next assessed behavior including locomotor activity and circadian rhythmicity in *Drosophila* chronically exposed to NPs (Fig.5A). Locomotor behavior was evaluated using the negative geotaxis assay, a standard method for detecting neurobehavioral deficits in flies. Chronic dietary NPs exposure at 0.01 mg/mL and 0.05 mg/mL significantly impaired climbing performance, with 24% and 39% of flies, respectively, failing to ascend 15 cm within 30 seconds compared with controls (Fig. 5B; P < 0.001). Statistical analysis confirmed significant locomotor deficits at both concentrations (P < 0.001). Actogram analysis revealed disrupted circadian organization, with reduced locomotor activity during the light phase and altered suppression of activity before dark onset, particularly in the 0.05 mg/mL group (Fig. 5C). These results indicate that chronic NPs exposure reduces overall activity and leads to progressive impairment of locomotor performance in a dose-dependent manner. Given that circadian rhythm impairments are frequently associated with mitochondrial dysfunction and dopaminergic deficits in PD [19], we next monitored daily activity–rest cycles under 12:12 h light–dark conditions for seven days. NPs exposure caused a dose-dependent reduction in overall daily activity, with a ∼2.5-fold decrease at 0.01 mg/mL (*P* = 0.002) and a ∼3.5-fold decrease at 0.05 mg/mL (*P* = 0.001; Fig. 5D & E). These findings indicate that NPs impair both amplitude and phase regulation of circadian rhythms, likely reflecting altered behaviour. This observation is consistent with earlier reports showing that NPs exposure can interfere with behavioral regulation in both mammalian models such as mice and invertebrate models such as *Drosophila* [23,72]. Collectively, these results demonstrate that chronic NPs exposure impairs locomotor function, disrupts circadian rhythm regulation, in *Drosophila*. These phenotypic outcomes, together with the observed neurochemical alterations, recapitulate core hallmarks of PD, reinforcing the notion that NPs may act as environmental stressors contributing to PD-like neurodegenerative processes.

**Figure 5:**
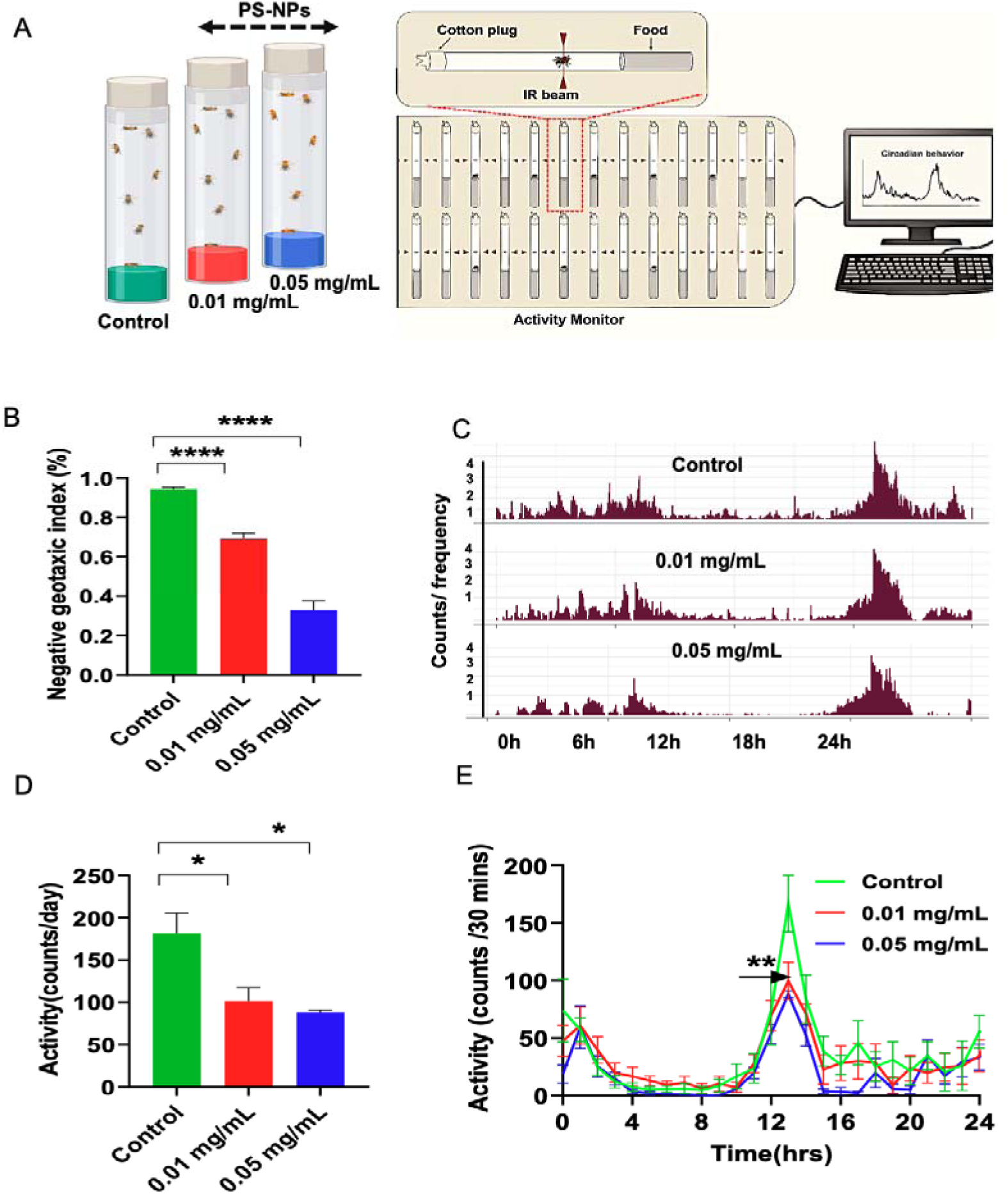
Behavioral disruption in *Drosophila* upon NPs exposure. **(A).** Experimental plan for PS-NPs treatment of flies**, (B)** Negative geotaxis activity bar plot showing significantly reduced movement in NPs-treated flies compared to control **(C)** 24-hour actograms of locomotor activity showing strong circadian rhythmicity in control flies, with disrupted rhythmic patterns in NPs-treated groups**. (D)** Activity per day index bar plot shows dose-dependent reduction in activity strength. **(F)**Time-series locomotor activity plot (counts per 30 min) reveals reduced morning peak activity in treated groups; significant difference observed at Zeitgeber time ZT13. Values are expressed as Mean ± Standard Error of Mean (n = 6). Significant differences from the control are indicated *p* <* 0.05, *p**<*0.005, *p****<*0.0001, student T-Test.

### 3.6 Antioxidant NAC confers protection against NPs-induced lipid metabolism in *Drosophila* brain

Our lipidomic profiling revealed significant NPs-induced disruptions in brain lipid homeostasis, characterized by enhanced neutral lipid accumulation and mitochondrial perturbations. ROS-mediated damage is recognized as a central mechanism of NPs-induced toxicity, implicating oxidative stress in lipid metabolism dysregulation, lipid droplet accumulation, lipid peroxidation and subsequent MUFAs accumulation. ROS-mediated damage is recognized as a central mechanism of NPs-induced toxicity, implicating oxidative stress in lipid metabolism dysregulation [73]. Since oxidative stress emerged as a key driver of these metabolic shifts, we next tested whether antioxidant supplementation could mitigate this phenotype. Flies were co-treated with NAC, (1 mg/ML for 7 days), a thiol antioxidant known to scavenge ROS, replenish intracellular glutathione, and preserve mitochondrial redox balance. Remarkably, NAC treatment led to a significant reduction in 0.05 mg/mL NPs-induced lipid droplet accumulation and lipid peroxidation in the *Drosophila* brain, restoring levels closer to those observed in controls (Fig.6A–C). Earlier studies demonstrating that NAC attenuates lipid droplet formation under oxidative stress conditions by preventing ROS-driven lipid peroxidation in PD disease models even in human studies [74]. For instance, in rotenone-based PD models, NAC supplementation has been shown to limit abnormal lipid storage. More recent reports highlight NAC’s capacity to counteract NPs-induced neuronal toxicity, including synaptic dysfunction and ROS-mediated injury signaling [11]. Our observations demonstrating that NAC not only mitigates oxidative damage but also restores lipid metabolic balance in NPs-exposed *Drosophila* brains. Lipidomic profiling further revealed that supplementation with NAC significantly reduced levels of the MUFAs oleic acid (C18:1) and palmitoleic acid (C16:1) in NPs-treated *Drosophila* brains (Fig.6D & E). This reduction is noteworthy, as it was previously reported identified abnormal accumulation of MUFAs as a hallmark of α-synuclein-induced neurotoxicity, mediated through dysregulation of stearoyl-CoA desaturase (SCD) activity. By attenuating MUFA levels, NAC may exert a protective effect against NPs-induced neurodegeneration, potentially through the suppression of lipid peroxidation substrates that otherwise promote oxidative stress, aggravate α-synuclein aggregation, and impair dopaminergic function. Importantly, we also observed a significant reduction in lipid peroxidation in NAC co-treated fly brains compared with NPs exposure alone, directly supporting the idea that normalization of MUFA levels reduces the availability of oxidizable substrates. Thus, the observed decrease in oleic and palmitoleic acid following NAC treatment suggests that the antioxidant not only scavenges reactive oxygen species but may also normalize lipid metabolic fluxes implicated in PD pathology. This aligns with previous reports that reducing MUFA availability can ameliorate α-synuclein toxicity in experimental models, thereby positioning NAC as a dual-acting neuroprotective agent in redox-lipid crosstalk [59].

**Figure 6.**
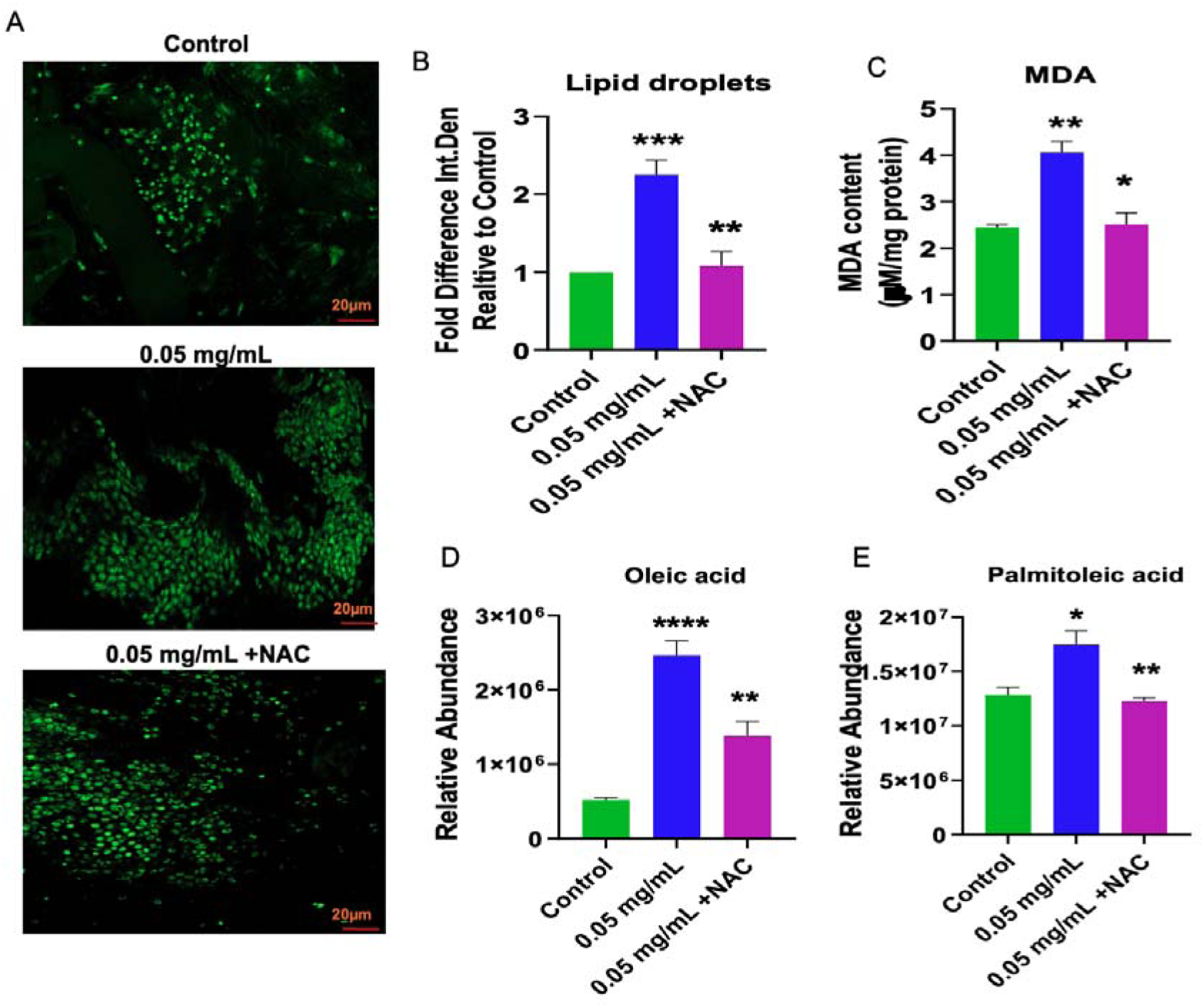
NAC mitigates NPs-induced lipid droplet accumulation, lipid peroxidation, and fatty acid imbalance in *Drosophila* brains. **(A)** Confocal images of BODIPY stained brains showing lipid droplet (LD) accumulation. LDs were markedly increased in NPs treated groups (0.05 mg/ML) but NAC co-treatment reduced LD density and clustering, restoring near-normal levels. Scale bar = 20 µm**. (B)** LD fold change significantly increased with NPs and was rescued by NAC. **(C**) Malondialdehyde (MDA) levels, elevated in NPs-exposed brains, were significantly reduced by NAC, indicating decreased lipid peroxidation. **(D–E)** Fatty acid profiling showed dysregulation of **(D)** Oleic acid (C18:1) **(E)** Palmitoleic acid (C16:1) in NPs-induced increases in these FAs were partially or fully normalized by NAC, suggesting restoration of lipid homeostasis. Values are expressed as Mean ± Standard Error of Mean (n = 5). Significant differences from the control are indicated p* < 0.05, p**<0.005, p****<0.0001 Student t-test.

### 3.7 NAC preserves mitochondrial function and TCA cycle integrity under NPs-induced stress

Because mitochondrial dysfunction is an early event in cellular stress responses, with ROS production driving downstream lipid peroxidation, lipid droplet accumulation often arises as a compensatory response to mitochondrial oxidative stress [75]. Conversely, excessive neutral lipid buildup can feed back to exacerbate mitochondrial impairment, creating a vicious cycle of redox imbalance and metabolic disruption. Building on our lipidomic observations of enhanced neutral lipid accumulation in NPs-exposed brains, we reasoned that mitochondria are a key subcellular target underlying this phenotype. Given the strong reciprocal link between lipid remodeling and mitochondrial impairment, we next examined whether NAC supplementation could protect mitochondrial function in NPs-exposed *Drosophila* brains. NAC co-treatment significantly preserved MMP compared with NPs-exposed flies, as indicated by restoration of MMP values toward control levels (Fig. 7A & B). In parallel, mitoSOX fluorescence revealed that NAC markedly reduced mitochondrial superoxide accumulation, suggesting effective scavenging of ROS within the organelle (Fig. 7C & D). NAC lowered total ROS (Fig 7E & F), and stabilized membrane potential. To assess whether these improvements extended to electron transport chain functionality, we measured the enzymatic activities of complex I and complex IV, which represent key entry and terminal points of the respiratory chain, respectively. NPs exposure led to substantial inhibition of both complexes, reflecting impaired electron transfer and increased vulnerability to oxidative stress. Strikingly, NAC co-treatment restored complex I and IV activities close to baseline levels (Fig. 7G & H), underscoring its capacity to protect mitochondrial bioenergetics. These findings align with earlier studies showing that NAC preserves mitochondrial structure and function in toxin-induced models of neurodegeneration, in part by replenishing glutathione and reducing oxidative stress.

**Figure 7.**
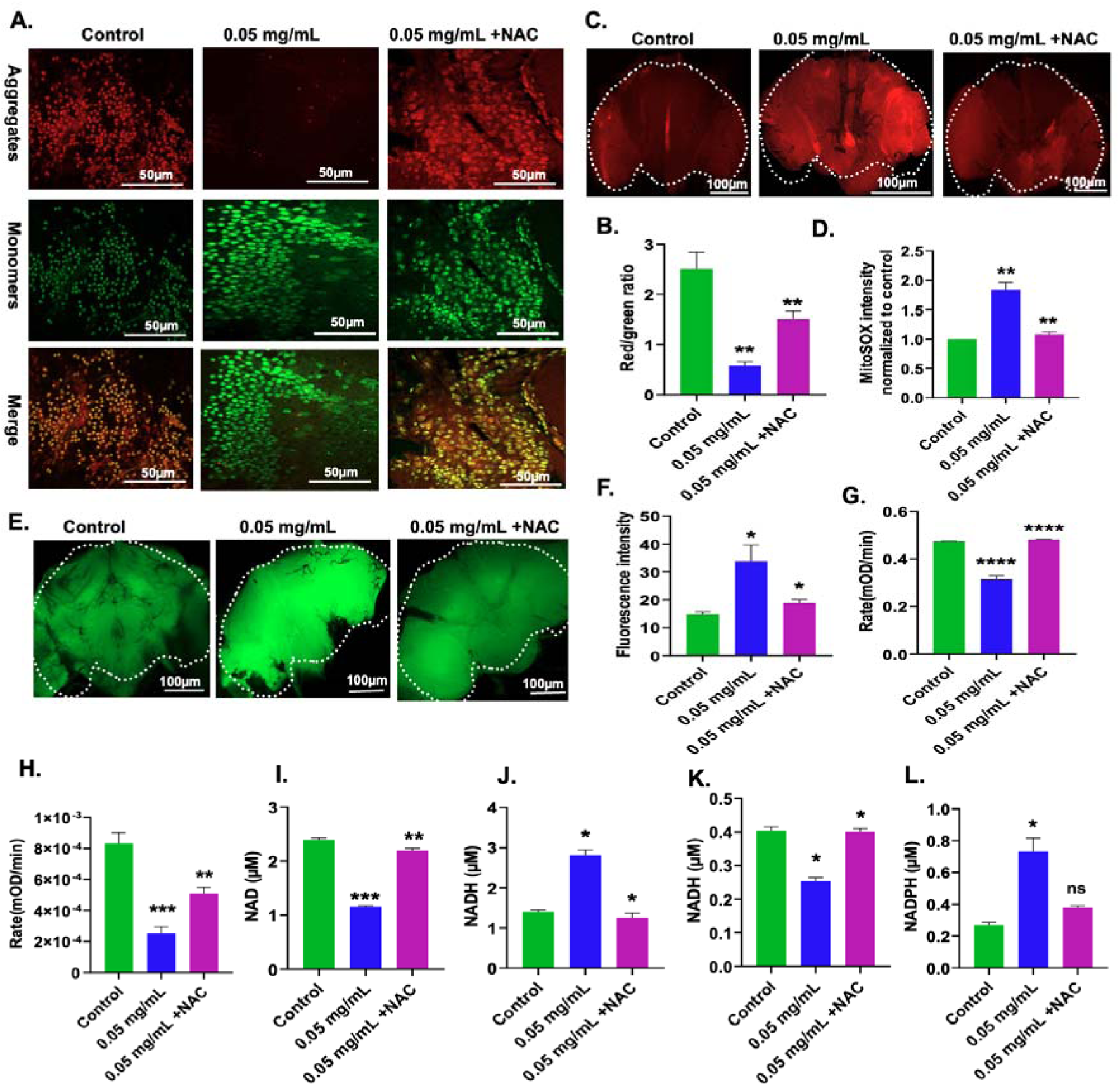
NAC mitigates NPs-induced mitochondrial dysfunction in *Drosophila* brains **(A)** Measurement of mitochondrial membrane potential (MMP) in brain tissue under the exposure of NPs with NAC. Scale bar = 50 µm (**B).** Bar graph showing the polarization state of MMP with red (polarization)and green (Depolarization) ratio. (**C)** Mitochondrial superoxide in brain tissue under the exposure of NPs with NAC. (**D)** MitoSOX intensity of mitochondrial superoxide in *Drosophila.* Brain Scale bar = 100 µm. **(E&F)** ROS intensity confocal images of 0.05 mg/mL NPs-treated brain tissue with NAC. Scale bar = 100 µm **(G)** NAC improved the mitochondrial complex 1activity. (**H)** NAC improved the mitochondrial Cytochrome C oxidase activity. **(I&J)** NAC improved the mitochondrial NAD and NADH concentration. (**K&L)** NAC improved the mitochondrial NADP and NADPH concentration. Values are expressed as Mean ± Standard Error of Mean (n = 5). Significant differences from the control are indicated p* < 0.05, p**<0.005, p****<0.0001 Student t-test.

In our study, NAC not only restored complex I and IV activities but also normalized NAD□ and NADH (Fig 7 I & J), NADP□ and NADPH (Fig 7k & L). Together, these results establish NAC as a potent redox modulator capable of reversing NPs-induced mitochondrial dysfunction and protecting neuronal energy metabolism in *Drosophila* brains.

Because the TCA cycle is directly coupled to respiratory chain activity and provides reducing equivalents (NADH) essential for oxidative phosphorylation, disruption of complex I and IV strongly predicts metabolic imbalance at the level of TCA intermediates. To directly probe whether NPs exposure alters carbon flux through central metabolism, we employed Uniform¹³C-labelled glucose feeding to *Drosophila* and traced incorporation into TCA cycle metabolites in brain samples. NPs-treated brain showed decrease in the contents of citric acid, malic acid, succinic acid and alpha ketoglutarate evident by labelling pattern. These results are consistent with a previous study that found NPs-treated liver cells (BEAS-2B cells) exhibited decreased levels of citrate [63]. Furthermore, the accumulation of oleic acid content indicates increased fatty acid oxidation by redirecting the flux from TCA cycle to fatty acid biosynthesis. Treating *Drosophila* with NAC protects rerouting of flux from TCA cycle to fatty acid oxidation from NPs induced stress evident by maintaining of the levels of TCA cycle metabolites and decreased labelling of oleic acid (Fig 8).

**Figure 8:**
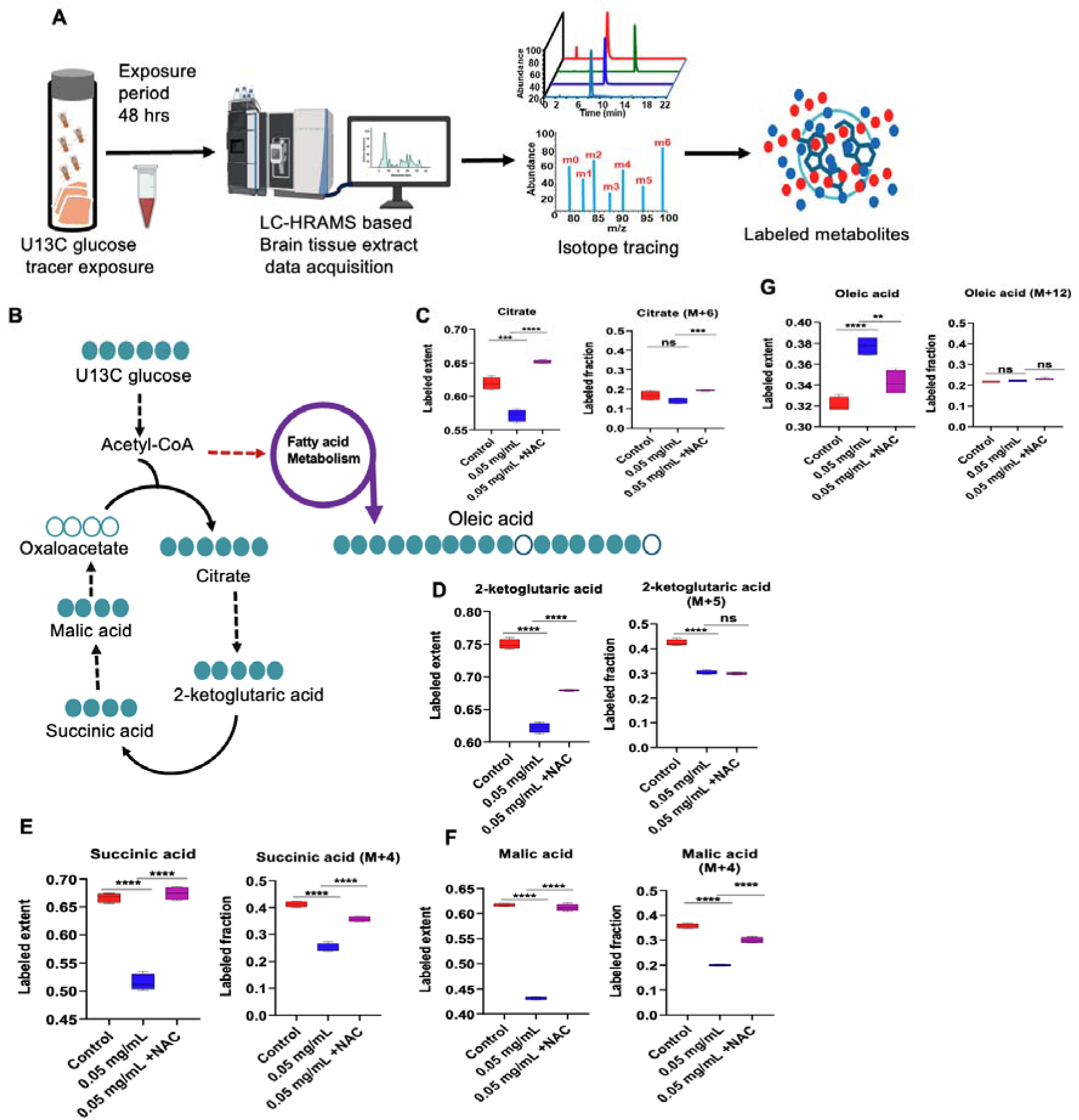
Metabolic dysregulation associated with TCA cycle under microplastic exposure in the brain tissue of *Drosophila. (***A).** Overview of the U13C glucose tracing framework using LC□MS/MS **(B).** Schematic illustrating the incorporation of carbon from U-13C-glucose into TCA cycle intermediate and fatty acid metabolism. (**C-F).** Box plots showing the labeling extent and labeling fraction of TCA cycle metabolites. (**G).** Box plots showing the labeling extent and labeling fraction of fatty acids metabolites. The significantly changed metabolites shown (\*\*\*\**p <*0.0001, two-tailed Student’s t-test).

Together with the observed decrease in MUFA accumulation and lipid peroxidation, our results highlight that NAC protects against NPs-induced neurotoxicity through dual mechanisms: normalization of redox–lipid interactions and stabilization of mitochondrial bioenergetics. By preserving both membrane lipid integrity and respiratory chain function, NAC may confer broad neuroprotective effects that counteract the converging pathways of oxidative stress, lipid metabolic dysregulation, and mitochondrial dysfunction.

### 3.8 NAC confers protection against NPs-induced pathological damage and improves behavior in *Drosophila* brain

Building on our earlier findings that redox–lipid metabolism plays a central role in NPs-induced neurotoxicity, and that NAC supplementation attenuates oxidative and metabolic stress, we next investigated whether NAC could protect against structural and neuronal alterations in the *Drosophila* brain. Scanning electron microscopy (SEM) analysis revealed that chronic NPs exposure caused profound ultrastructural damage. In control brains, the surface appeared smooth and continuous, with well-preserved morphology. By contrast, NPs-exposed brains exhibited significant surface deformation, including ruptures, irregular contours, and loss of structural integrity, particularly at higher magnifications (50 µm and 10 µm) (Fig. 9A). These morphological aberrations suggest compromised membrane stability and extracellular architecture, consistent with oxidative and lipid peroxidation-driven damage. Remarkably, co-treatment with NAC visibly ameliorated these defects. NAC-treated brains showed smoother surfaces, reduced deformation, and partial restoration of surface organization, indicating that antioxidant supplementation can preserve brain tissue integrity even under chronic NPs stress.

**Figure 9.**
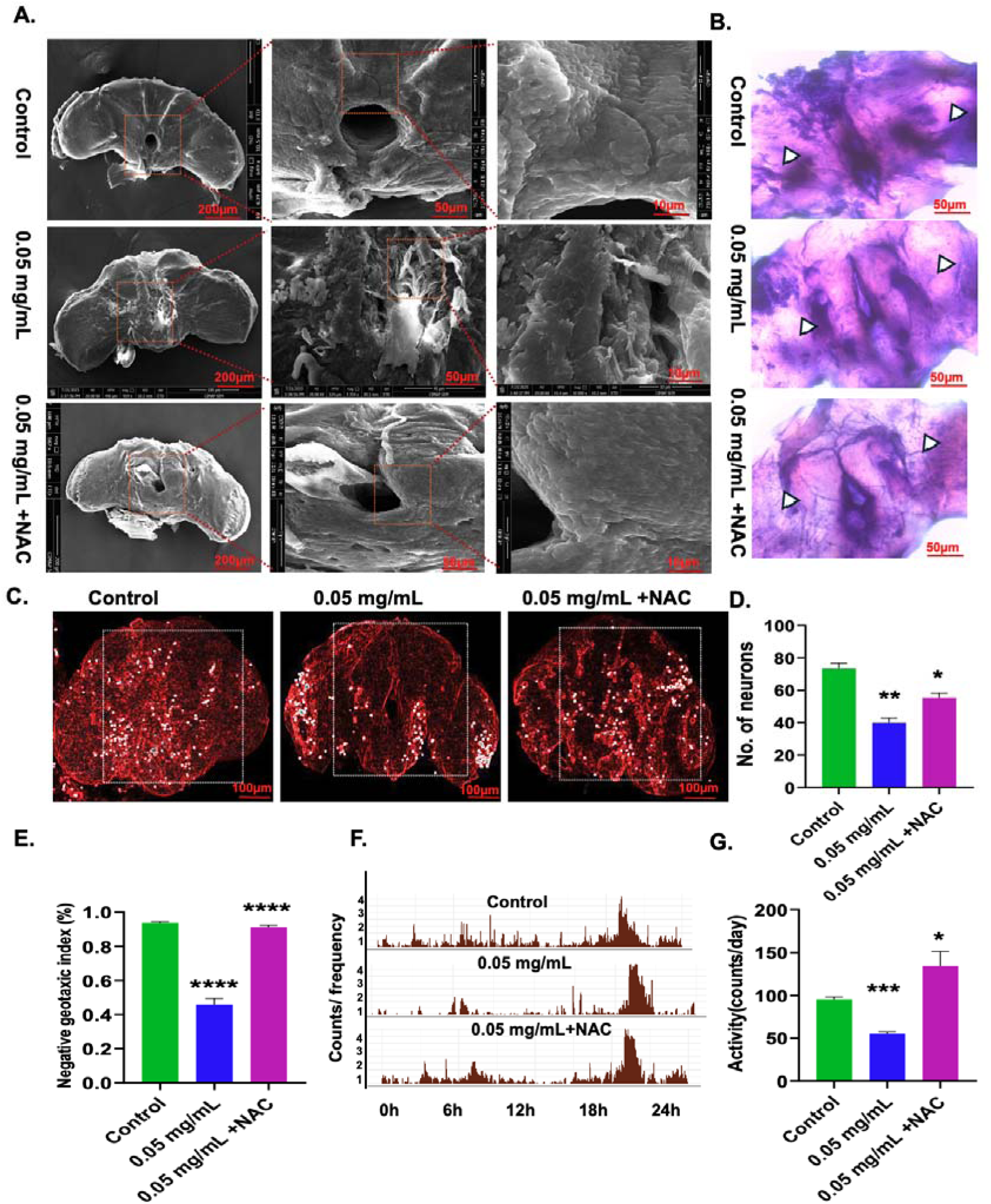
NAC improved the NPs-Induced structural damage. (A) SEM images of *Drosophila* brain at increasing magnifications (200 µm, 50 µm, and 10 µm) reveal surface formation NAC improved the damage in treated groups. (B) H&E-stained brain sections exhibit NAC decrease vacuolization (white arrowheads) in NPs-exposed brains. (C&D) Immunofluorescent staining of tyrosine hydroxylase (TH)-positive neurons in *Drosophila* brains, showing distribution and intensity differences across experimental groups. (E) Negative geotaxis activity bar plot showing NAC significantly improved movement in NPs-treated flies. (F) 24-hour actograms of locomotor activity showing strong circadian rhythmicity in control flies, with NAC improved rhythmic patterns in NPs-treated groups. (G) Activity per day index bar plot shows NAC improved in activity strength.

**Figure 10.**
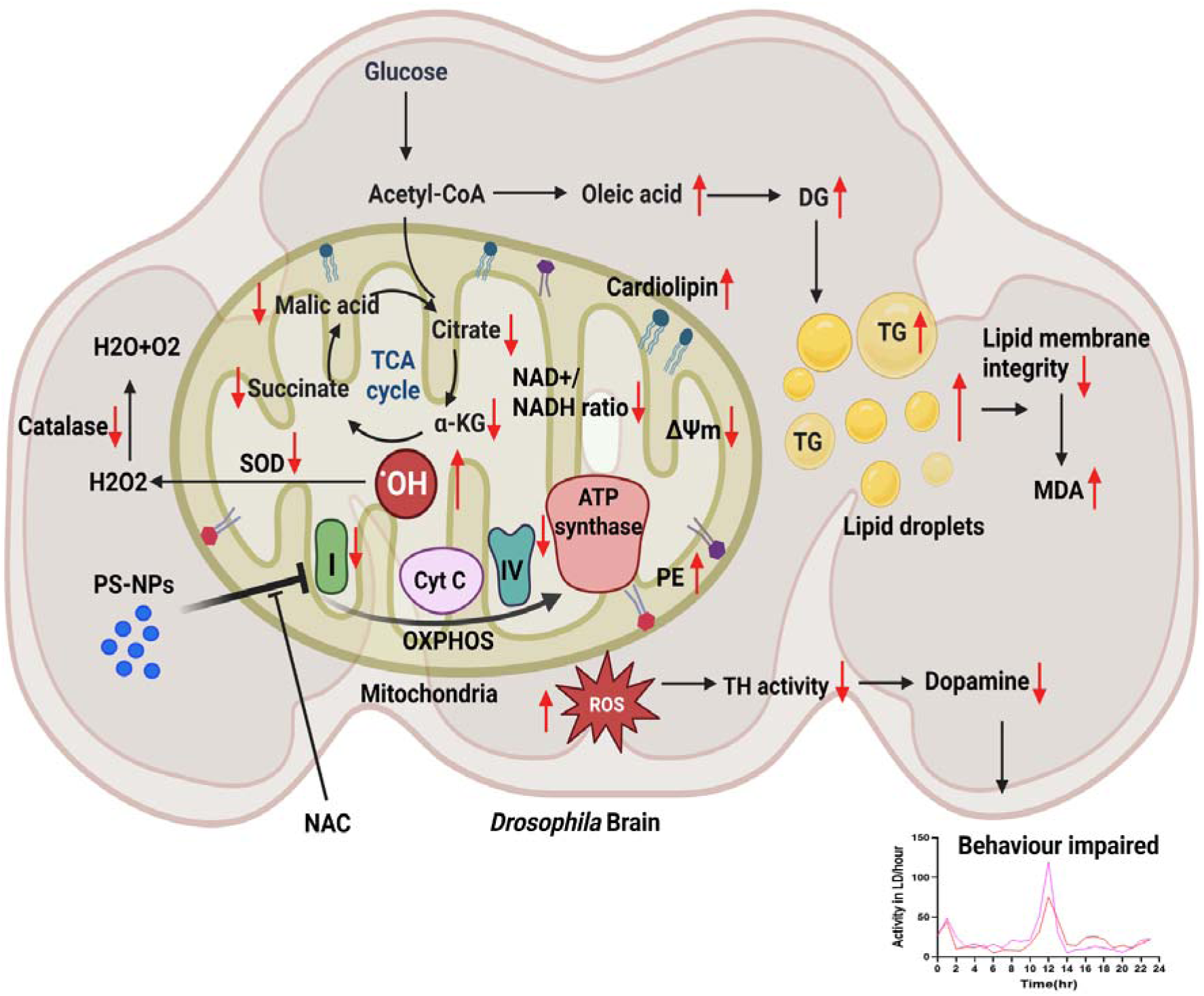
Proposed mechanistic model of PS-NP–induced neurotoxicity in the *Drosophila* brain. Exposure to PS-NPs leads to mitochondrial dysfunction and oxidative stress, disrupting brain energy metabolism and neurotransmission. PS-NPs impair Complex I–mediated oxidative phosphorylation (OXPHOS), reducing mitochondrial membrane potential (ΔΨm), and NAD□/NADH ratio. The resulting accumulation of ROS (•OH, H□O□) overwhelms antioxidant defenses (SOD, catalase), elevating MDA and compromising membrane integrity. Dysregulated TCA intermediates (citrate, α-ketoglutarate, succinate, malate) and altered phospholipids (cardiolipin, phosphatidylethanolamine) further destabilize mitochondrial structure. Excess oleic acid and DG promote TG accumulation in lipid droplets, reflecting disturbed lipid homeostasis. These biochemical imbalances suppress TH activity and dopamine synthesis, leading to behavioral impairment. Antioxidant supplementation with NAC partially mitigates ROS accumulation and preserves mitochondrial integrity.

Complementary histological evaluation using hematoxylin and eosin (H&E) staining further supported these observations. NPs-exposed brains displayed extensive vacuolization, disorganization of neuropil structure, and tissue rarefaction, as highlighted by white arrowheads (Fig. 9B). Such vacuolar degeneration is a well-recognized hallmark of neurotoxic injury, reflecting loss of neuronal processes and cellular swelling. Strikingly, NAC treatment substantially reduced vacuolization and preserved overall brain architecture, reinforcing its protective role against NPs-induced structural neurodegeneration.

To assess neuronal vulnerability more specifically, we examined dopaminergic neurons using immunofluorescence labeling of tyrosine hydroxylase (TH), the rate-limiting enzyme in dopamine biosynthesis and a critical marker of dopaminergic integrity. Control flies exhibited a well-organized distribution of TH-positive neuronal clusters with strong fluorescence intensity. In contrast, NPs exposure caused a marked reduction in TH immunoreactivity, accompanied by fragmentation of dopaminergic clusters and loss of neuronal projections (Fig. 9C & D). These effects were most pronounced in the protocerebral and subesophageal regions, which are known to regulate locomotion and feeding behavior, suggesting region-specific vulnerability of dopaminergic circuits to NPs-induced neurotoxicity. Notably, NAC co-treatment significantly preserved dopaminergic neuronal integrity. Brains from NPs+NAC groups displayed higher TH fluorescence intensity, better-organized neuronal clusters, and reduced fragmentation compared to NPs-only groups. NAC significantly attenuated neuronal loss and maintained TH expression, highlighting its neuroprotective action. These findings support the notion that NAC acts through redox modulation to counteract NPs-induced oxidative stress, thereby protecting vulnerable dopaminergic neurons. These results demonstrate that NAC supplementation confers structural and functional neuroprotection against NPs-induced brain pathology in *Drosophila.* By reducing vacuolization, preserving tissue architecture, and maintaining dopaminergic neuronal integrity, NAC mitigates both cellular and regional hallmarks of neurodegeneration, further underscoring the therapeutic potential of targeting redox–lipid pathways in NPs-associated neurotoxicity.

Having established that NAC preserves mitochondrial integrity, lipid homeostasis, and dopaminergic neuronal structure under NPs stress, we next asked whether these protective effects extend to higher-order outcomes such as locomotor performance and circadian rhythmicity. Supplementation with NAC markedly rescued these behavioral impairments. In the negative geotaxis assay, NAC-treated flies maintained climbing ability comparable to controls, counteracting the dose-dependent decline observed under NPs exposure (Fig. 9E). Similarly, NAC preserved circadian rhythmicity, preventing both the reduction in daily activity and the phase misalignment induced by NPs (Fig. 9F & G). These findings demonstrate that NAC not only protects cellular and neuronal integrity but also sustains behavioral resilience, thereby mitigating NPs-induced neurotoxicity in vivo.

### 3.9 Correlation of NPs-induced alterations in *D. melanogaster* with Human PD Pathology

More importantly, we have presented significant NPs-induced changes in this study, including altered lipid metabolites and multiple biochemical parameters, that closely correlate with those observed in postmortem brain samples from human PD subjects (Table 1). The parallel upregulation of key lipid classes, including DG, TG, PE, and cardiolipin, along with increased lipid droplet accumulation and oleic acid dysregulation, highlights substantial conservation of lipid metabolic vulnerability between the fly model and human PD brains. Likewise, NPs induced impairments in locomotor performance, circadian rhythmicity, dopamine and GABA levels, TH activity, and mitochondrial function strongly align with hallmark neurochemical and behavioral deficits in PD. The consistent elevation of oxidative stress markers (ROS, mitochondrial ROS, MDA) and suppression of antioxidant defenses (catalase, SOD), together with disruptions in NAD(H)/NADP(H) homeostasis and TCA cycle metabolites, further underscores a shared metabolic and bioenergetic collapse characteristic of PD pathophysiology. Collectively, these converging alterations reinforce *D. melanogaster* as a robust model for investigating NPs-induced neurotoxicity and suggest that NPs exposure may recapitulate key biochemical and mitochondrial dysfunctions implicated in human Parkinson’s disease.

**Table 1:** Correlation of NPs induced altered lipid metabolites with human PD samples.

| S.NO | Altered Metabolism/ Important biological parameters | Human-PD brain | NPs induced *D. melanogaster* brain | References<sup>a</sup> |
| --- | --- | --- | --- | --- |
| 1. | DG lipid metabolism | Upregulation | Upregulation | [S1] |
| 2. | TG lipid metabolism | Upregulation | Upregulation | [S2] |
| 3. | Phosphatidylethanolamine (PE) lipids metabolism | Upregulation | Upregulation | [S3] |
| 4. | Cardiolipin (CL) metabolism | Upregulation | Upregulation | [S4] |
| 5. | Fatty acids | Oleic acid in neuronal cells | Oleic acid in brain upregulates | [S5] |
| 6. | Lipid droplets | Upregulation | Upregulation | [S6] |
| 7. | Altered Locomotor activity | Decrease the locomotor activity | Decrease the locomotor activity | [S7] |
| 8. | Circadian rhythms | Decrease in amplitude with increased day sleep | Decrease in amplitude with increased day sleep | [S8] |
| 9. | Dopamine level | Downregulation | Downregulation | [S9, S10] |
| 10. | GABA level |  | Downregulation | [S11] |
| 11. | TH activity/ dopaminergic system | Downregulation | Downregulation | [S12] |
| 12. | ROS level | Upregulation | Upregulation | [S13] |
| 13. | MDA level | Upregulation | Upregulation | [S14] |
| 14. | Catalase Activity | Downregulation | Downregulation | [S15] |
| 15. | SOD Activity | Downregulation | Downregulation | [S16] |
| 16. | Mitochondrial membrane potential | Downregulation | Downregulation | [S17] |
| 17. | Mitochondrial ROS | Upregulation | Upregulation | [S18] |
| 18. | Mitochondria Complex 1 | Downregulation | Downregulation | [S19] |
| 19. | Mitochondria Complex IV | Downregulation | Downregulation | [S20] |
| 20. | NAD(H) | Dysregulation | Dysregulation | [S21] |
| 21. | NADP(H) | Dysregulation | Dysregulation | [S22] |
| 22. | TCA cycle metabolites | Downregulation | Downregulation | [S23, S24] |

<sup>a</sup>References S1 to S24 are specified at supplementary information

## 4. Conclusions

This study, for the first time, provides a comprehensive understanding of how NPs exposure perturbs the brain lipidome and mitochondrial function using *Drosophila* as a model system. Using comprehensive lipidomics, we demonstrate that NPs accumulate in the brain, disrupt membrane integrity, and drive profound remodeling of lipid metabolism. Our findings reveal a dose-dependent elevation in DG and TG pools, accompanied by enhanced lipid droplet biogenesis and enrichment of MUFAs, hallmarks of oxidative stress–driven lipid remodeling. These alterations reflect an adaptive but ultimately maladaptive response, linking NPs exposure to disrupted membrane homeostasis and redox imbalance. Importantly, we identify mitochondrial dysfunction as a central event underlying these lipid perturbations. NPs exposure caused a marked loss of mitochondrial membrane potential, elevated mtROS production, and inhibition of respiratory chain complexes I and IV, highlighting mitochondria as sensitive subcellular targets of NPs toxicity. These mitochondrial impairments were closely associated with reduced levels of dopaminergic neurotransmitters and disrupted GABA signaling, thereby providing a mechanistic basis for the behavioral and survival deficits observed in NPs-exposed flies. Remarkably, antioxidant supplementation with NAC conferred broad neuroprotection. NAC not only reduced lipid droplet accumulation and normalized MUFA levels but also preserved mitochondrial membrane potential, decreased mitochondrial ROS, and restored the activities of respiratory chain complexes. Furthermore, NAC attenuated structural brain damage, preserved dopaminergic neuronal integrity, and improved locomotor and circadian behaviors. Together, our findings establish a clear mechanistic framework in which NPs exposure perturbs redox–lipid interactions and mitochondrial bioenergetics, thereby driving neurodegenerative phenotypes. By demonstrating that NAC mitigates these converging pathways, this study provides novel insight into the therapeutic potential of targeting oxidative stress and lipid metabolic remodeling in NPs-induced neurotoxicity.

## CRediT authorship contribution statement

Ratnasekhar CH: Conceptualization, Formal analysis, Funding acquisition, Investigation, Project administration, Software, Supervision, Writing – original draft, – review and editing. Priya Rathor: Methodology, Data curation, Formal analysis, Validation, Visualization, Writing - review & editing. Ashutosh Kumar Tiwari: Methodology, Data curation, Formal analysis, Validation, Visualization, Writing - review & editing. Rajendra Patel: Data curation, Formal analysis, Visualization, Writing - review & editing. Sheelendra Pratap Singh Data curation, Formal analysis, Writing - review & editing. Nick Birse: Resources, Methodology, Writing - review & editing.

## Acknowledgements

Ratnasekhar CH acknowledges funding support from the Science and Engineering Research Board, DST (Grant No. SRG/2021/000750-G), and from the Department of Biotechnology, Government of India, through the Ramalingaswami Re-entry Fellowship (BT-RLF-Reentry-21-2020). We gratefully acknowledge the High-Throughput Instrumentation Facility at CSIR-CIMAP for providing analytical support. Priya Rathor received funding from a UGC-SRF fellowship, Ashutosh Kumar Tiwari was supported by a CSIR Fellowship. We sincerely thank the Director, CSIR-CIMAP, for institutional facilities and infrastructure.

## Supplementary Data

Supplementary information consisting of Fig.S1-Fig S6 is provided.

## Data availability

Lipidomics data are submitted on the Metabolights Platform https://www.ebi.ac.uk/metabolights/editor/REQ20251016213928/assays

## Abbreviations

PS: Polystyerene
NPs: Nanoplastics
MPs: Microplastics
NAC: N-acetyl cysteine
BBB: Blood-brain barrier
PD: Parkinson’s Disease
HR-MS: High-resolution mass spectrometry
PE: Phosphatidylethanolamine
CL: Cardiolipin
DG: Diglyceride
TG: Triglyceride
ROS: Reactive oxygen species
HEPES: 4-(2-Hydroxyethyl) piperazine-1-ethanesulfonic acid
SDS: Sodium dodecyl sulfate
TBA: Thiobarbituric acid
TEP: Tetraethoxypropane
DCFH-DA: Dichloro-dihydro-fluorescein diacetate
NBT: Nitroblue tetrazolium
TEM: Transmission electron microscopy
SEM: Scanning electron microscopy
DLS: Dynamic light scattering
PFA: Paraformaldehyde
TBARS: Thiobarbituric acid reactive substances
PMS: Phenazine methosulfate
H&E: Hematoxylin and eosin
TH: Tyrosine hydroxylase
FA: Fatty acyls
GL: Glycerolipids
GP: Glycerophospholipids
SL: Sphingolipids
ST: Sterol lipids
PLS-DA: Partial least squares discriminant analysis
MUFAs: Monounsaturated fatty acids
ER: Endoplasmic reticulum
mROS: Mitochondrial reactive oxygen species
SCD: Stearoyl-CoA desaturase

