## Supplementary information for "Brain lipidomics identifies mitochondrial redox dysfunction and metabolic trade-offs associated with Parkinson’s disease-like pathology induced by Nanoplastics exposure"

*Corresponding author

**ORCID ID:** [**0000-0001-7041-6827**](https://orcid.org/0000-0001-7041-6827)

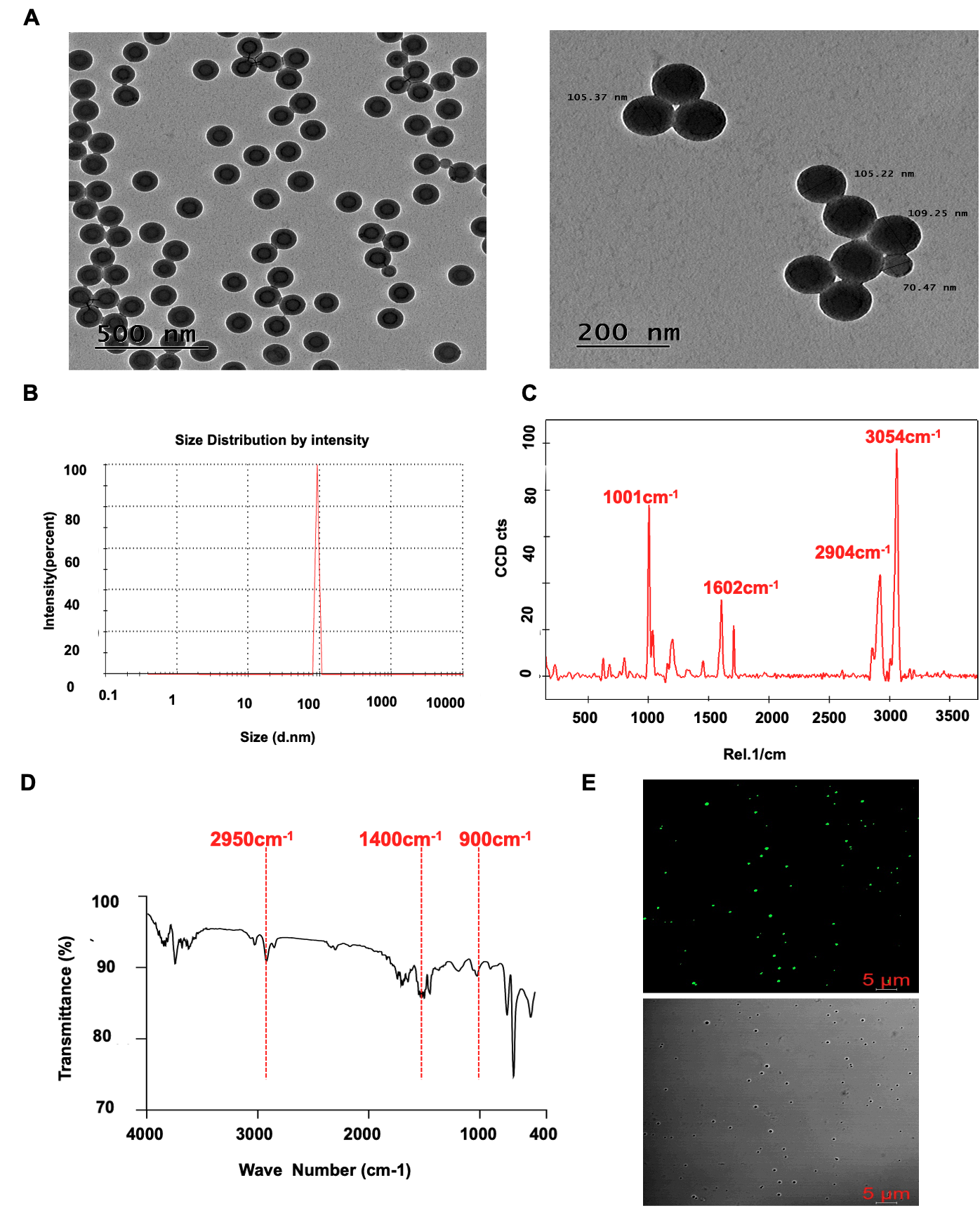

**Fig. S1**: Chemical characterization, of PS-NPs*.* (**A)** TEM image confirming particle morphology and uniformity with an average size distribution of ~100 μm. (**B)** DLS analysis showing narrow particle size distribution and good dispersion stability in aqueous media. **(C)** Raman spectra of PS-NPs validating chemical identity via prominent fingerprint peaks associated with aromatic polystyrene structure. **(D)** FT-IR spectra of synthesized PS-NPs showing characteristic vibrational peaks at ~900 cm⁻¹ (C–H aromatic bending), ~1400 cm⁻¹ (C=C aromatic stretching), and ~2950 cm⁻¹ (C–H stretching). **(E**) Confocal image of fluorescent labelled NPs.

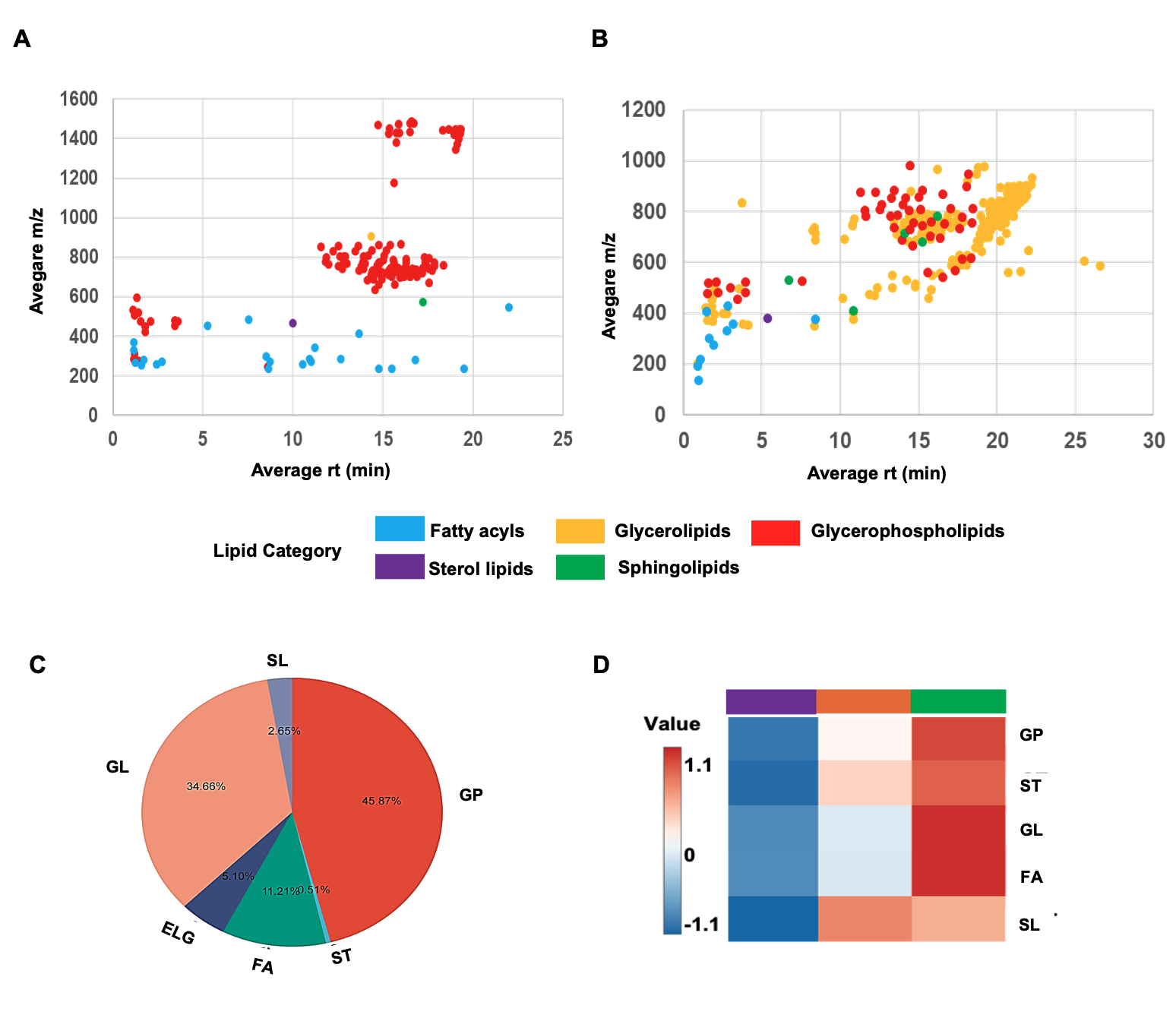

**Fig. S2**: Scatter plots showing the average m/z and average rt of detected lipids in brain tissue under the microplastic exposure (**A)** ESI Negative (**B)** ESI Positive. **(C)** Total lipid species present in *Drosophila* Brain tissue. **(D)** Lipid category response to NPs exposure in *Drosophila* brain tissue, fold change >1.5.

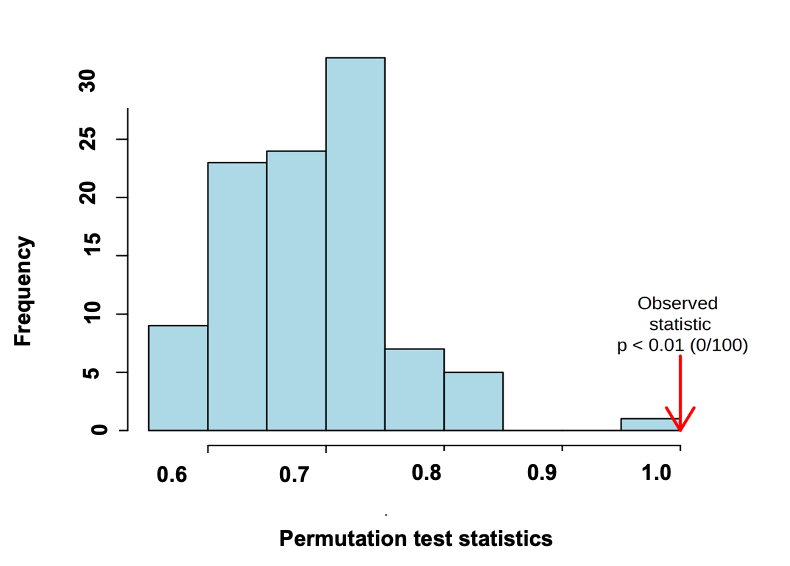

**Fig S3:** The histogram represents 100 permuted values of the test statistic, while the red arrow indicates the observed statistic from the actual data. Since none of the permuted values exceeded the observed value (0/100), the result is statistically significant with p < 0.01, suggesting that the observed effect is unlikely to have occurred by chance.

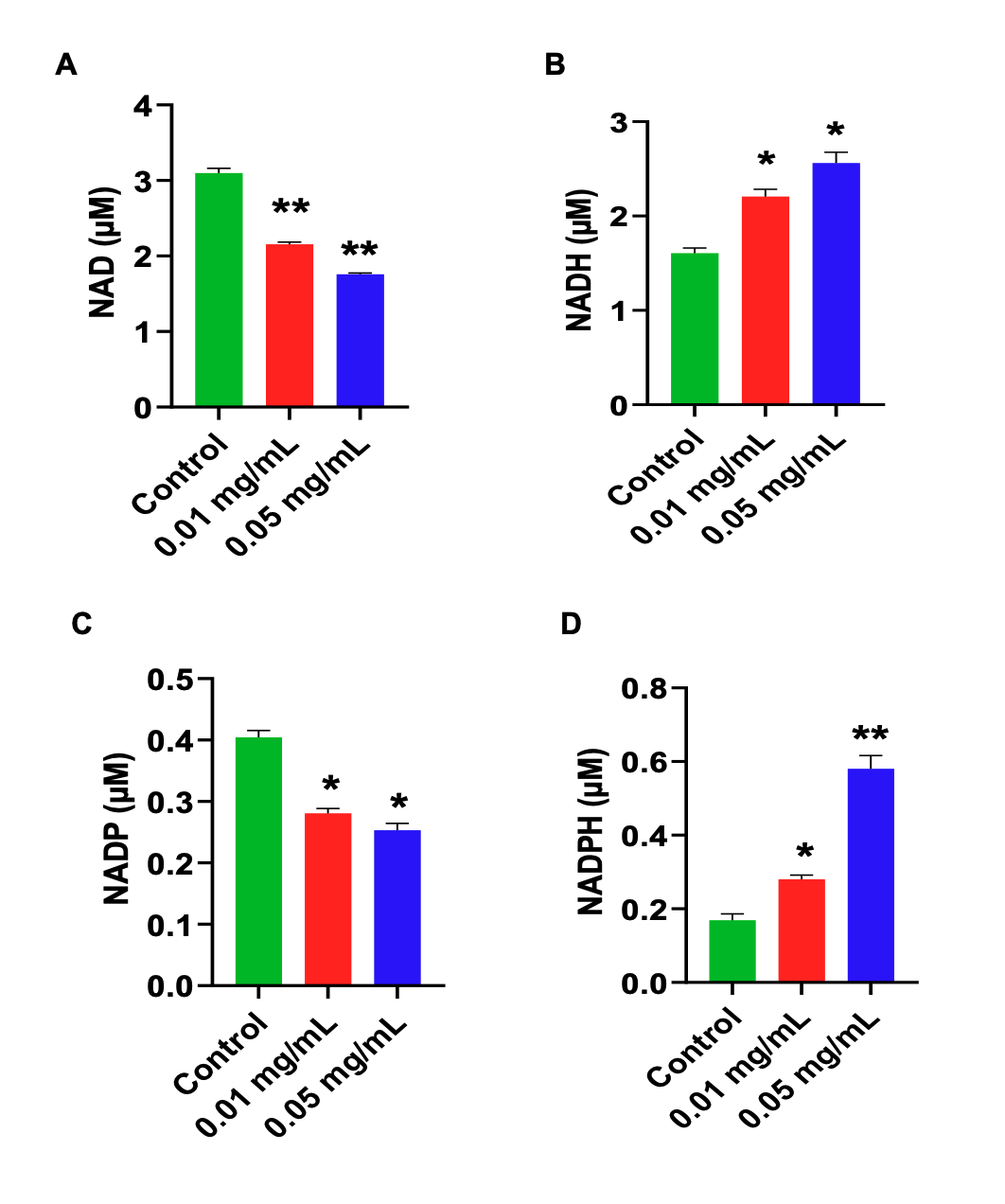

**Fig S4**: Effect of NPs exposure on redox cofactors at concentrations0.01, and 0.05 mg/ML in *Drosophila* brain. Bar graphs show mean ± SEM of A) NAD⁺, B) NADH, C) NADP⁺ D) NADPH. Values are expressed as Mean ± Standard Error of Mean (n = 5). Significant differences from the control are indicated p* < 0.05, p**<0.005, p****<0.0001 Student t-test.

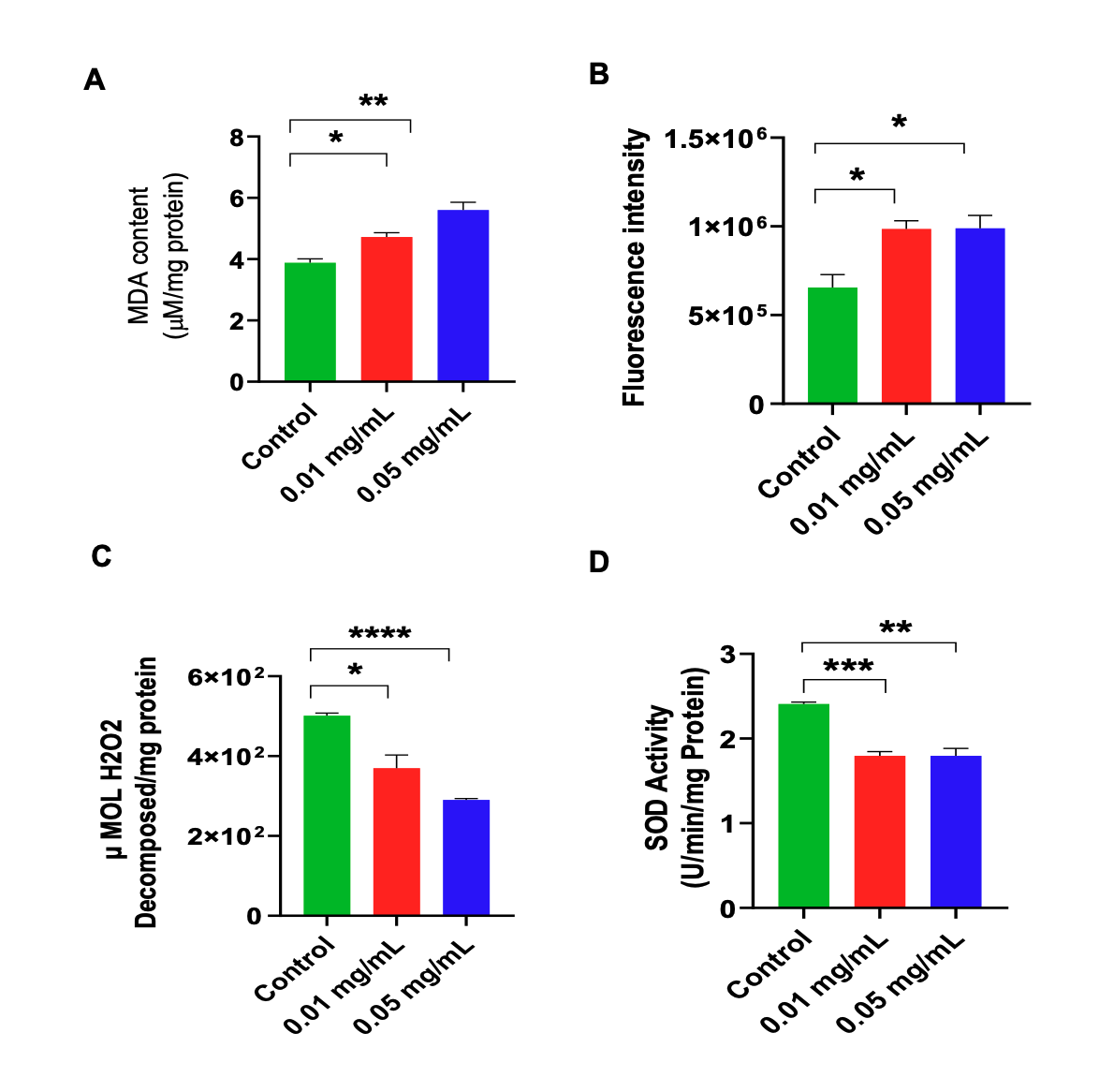

**Fig S5. A)** Elevated MDA, B) ROS and reduction in C) catalase, D) SOD activities, indicating oxidative stress and impaired mitochondrial redox balance and altered metabolic flux under PS-NP-induced oxidative stress. Values are expressed as Mean ± Standard Error of Mean (n = 5). Significant differences from the control are indicated p* < 0.05, p**<0.005, p****<0.0001 Student t-test.

| Table 1- Identified Marker lipids in fly brain tissue under the PS-NPs | | | | | | | | |
| --- | --- | --- | --- | --- | --- | --- | --- | --- |
| **Lipid Species** | **Lipid ID** | **Molecular formula** | **RT (min)** | **Adduct** | **Theoretical m/z** | **Observed m/z** | **ppm error** | **MS/MS** |
| CL 82:1 | FDB097114 | C91H176O17P2 | 17.3 | [M-H]- | 1603.3012 | 1603.3012 | 0 | 119.035, 256.2370,392.3891,593.5713, 640.3049,755.6237 |
| CL 66:1 | CID118701802 | C75H144O17P2 | 15.75 | [M-H]- | 1377.9806 | 1377.9913 | 7.76 | 229.0979, 255.2329, 661.5193,688.4918,689.4951,715.5116 |
| CL 70:3 | CID118702304 | C79H148O17P2 | 15.92 | [M-H]- | 1430.0119 | 1430.023 | 7.76 | 229.0660, 279.2327,689.4943, 730.30396,741.5261, 1366.5433 |
| CL 78:3 | SLM:000550285 | C87H164O17P2 | 16.28 | [M-H]- | 1543.1631 | 1543.1631 | 0 | 255.2329, 277.2172,281.2490, 392.3518,725.5766, 770.5624 |
| CL 70:5 | CID118702555 | C79H144O17P2 | 15.3 | [M-H]- | 1425.9806 | 1425.9917 | 7.78 | 229.1614, 255.2329,281.2490, 1117.3786, 1406.3568,1425.8072 |
| PE 32:3 | CID75957888 | C37H68NO8P | 14.15 | [M-H]- | 684.461 | 684.4611 | 0.14 | 203.0865, 229.0939, 424.246, 659.1667, 684.4603 |
| PE 28:0 | CID9852308 | C33H66NO8P | 14.57 | [M-H]- | 634.4454 | 634.4453 | -0.15 | 229.0831, 255.2332,289.1455, 424.2468,478.4002, 634.4425 |
| PE 36:4 | CID9546800 | C41H74NO8P | 15.37 | [M-H]- | 738.508 | 738.5081 | 0.13 | 203.0860,229.0965,279.2329, 477.2824, 738.5079 |
| PE 34:4 | CID52924148 | C39H70NO8P | 14.35 | [M-H]- | 710.4767 | 710.4768 | 0.14 | 278.2206, 279.2333, 450.2624,669.1933,710.4764 |
| PE 30:0 | CID20843280 | C35H70NO8P | 15.64 | [M-H]- | 662.4767 | 662.4769 | 0.3 | 227.2016, 229.7451, 255.2329,424.2471, 662.4724 |
| PE 34:1;O | CID52926191 | C39H76NO9P | 14.42 | [M-H]- | 732.5185 | 732.5272 | 11.87 | 255.2488,281.2481, 291.6192, 506.4570, 687.5246,732.4821 |
| PE 30:1 | CID52924132 | C35H68NO8P | 14.76 | [M-H]- | 660.461 | 660.461 | 0 | 229.1019, 255.2344, 450.2631, 660.4614 |
| PE 34:3 | CID9546746 | C39H72NO8P | 15.31 | [M-H]- | 712.4923 | 712.49231 | 0.01 | 255.2330, 280.2366, 452.2778,495.7064, 712.4923 |
| PE 32:2 | CID90657244 | C37H70NO8P | 14.94 | [M-H]- | 686.4767 | 686.4766 | -0.14 | 253.2171,287.3793, 340.5578:7089 ,450.2619,686.4758 |
| DG 42:10 | CID9543945 | C45H68O5 | 13.81 | [M+Na]+ | 711.4958 | 711.4941 | -2.38 | 229.1220,275.6198, 311.2572, 451.4545,711.4931 |
| DG 32:1 | CID9543678 | C35H66O5 | 17.12 | [M+Na]+ | 589.4802 | 589.4807 | 0.84 | 229.1144,291.3054, 333.2400, 433.4001, 589.5551 |
| DG 33:0 | CID131800229 | C36H70O5 | 25.66 | [M+Na]+ | 605.5115 | 605.5137 | 3.63 | 271.4818, 293.1744, 461.3993, 587.5045,605.5111 |
| DG 46:9 | CID129882426 | C49H78O5 | 10.95 | [M+Na]+ | 769.5741 | 769.5735 | -0.77 | 381.2996, 391.9769, 502.3153, 751.6030, 769.5736 |
| DG 34:1 | CID5282283 | C37H70O5 | 17.78 | [M+Na]+ | 617.5115 | 617.5117 | 0.32 | 263.2375,305.2443, 313.2953,361.2714, 575.4304,617.5858 |
| DG 30:0 | CID13734178 | C33H64O5 | 16.04 | [M+Na]+ | 563.4645 | 563.4668 | 4.08 | 277.5116, 294.1782,335.2205, 461.3991,563.4669 |
| DG 34:2 | CID9543695 | C37H68O5 | 17.17 | [M+NH4]+ | 610.5404 | 610.541 | 0.98 | 384.1442, 477.3924, 593.3606,609.6004, 610.6467 |
| DG 32:1 | CID9543678 | C35H66O5 | 17.09 | [M+NH4]+ | 584.5248 | 584.5241 | -1.19 | 277.1702, 285.2427, 311.2928,511.3747, 584.5620 |
| DG 44:6 | CID131801967 | C47H80O5 | 10.88 | [M+NH4]+ | 742.6343 | 742.6358 | 2.01 | 229.1128,285.2784,381.2966, 629.5307, 742.5701 |
| DG 30:1 | CID53477950 | C33H62O5 | 16.65 | [M+Na]+ | 561.4489 | 561.4511 | 3.91 | 291.1590,351.2171, 421.3320, 477.3935,561.4520 |
| TG 44:3 | CID131758749 | C47H84O6 | 19.49 | [M+NH4]+ | 762.6605 | 762.6602 | -0.39 | 272.3095,285.2416, 311.2582,519.4407, 546.4594,762.6635 |
| TG 48:3 | CID9543989 | C51H92O6 | 20.54 | [M+NH4]+ | 818.7231 | 818.7222 | -1.09 | 311.2577, 550.4913,575.5031, 801.6948,818.7210 |
| TG 38:1 | CID3034361 | C41H76O6 | 18.79 | [M+NH4]+ | 682.5979 | 682.5977 | -0.29 | 285.2435,327.0778, 411.3456,467.4094, 552.3327,682.5982 |
| TG 54:7 | CID14325545 | C57H96O6 | 20.27 | [M+NH4]+ | 894.7544 | 894.754 | -0.44 | 203.0856, 280.2612, 495.4383, 597.4873,600.5065, 894.8668 |
| TG 42:2 | CID6438769 | C45H82O6 | 19.44 | [M+Na]+ | 741.6003 | 741.6 | -0.4 | 311.2573,515.4074,541.4218, 601.5183, 741.5997 |
| TG 46:2 | CID56936614 | C49H90O6 | 20.51 | [M+Na]+ | 797.6629 | 797.6628 | -0.12 | 203.0856,229.1219,321.3064,569.4534, 797.6623 |
| TG 37:0 | CID131773487 | C40H76O6 | 18.95 | [M+NH4]+ | 670.5979 | 670.5979 | -0.12 | 271.2269, 445.2869,501.3491, 669.5938,670.6089 |
| TG 43:2 | CID56936607 | C46H84O6 | 20.48 | [M+NH4]+ | 750.6687 | 750.6657 | -3.9 | 229.1129,285.2427, 491.4106,533.4563, 750.8077 |
| TG 32:0 | CID99121155 | C35H66O6 | 17.37 | [M+NH4]+ | 600.5059 | 600.5195 | 5.99 | 233.1539, 355.2838,411.3468, 545.3830,600.5042 |
| TG 55:4;O2 | CID87527240 | C58H104O8 | 18.26 | [M+NH4]+ | 946.8151 | 946.807 | -8.55 | 285.2434,311.2764, 675.5556, 683.5606,947.2888 |
| TG 42:2 | CID6438769 | C45H82O6 | 19.44 | [M+Na]+ | 741.6003 | 741.6004 | 0.13 | 311.2573,411.2038, 487.3757, 741.5997 |
| TG 42:3 | CID14122758 | C45H80O6 | 19 | [M+NH4]+ | 734.6374 | 734.6292 | -11.16 | 283.2266,311.2587, 492.4127,520.4439, 734.6423 |
| TG 42:1 | CID131753215 | C45H84O6 | 19.8 | [M+Na]+ | 743.6159 | 743.6153 | -0.8 | 211.2057,229.1285,237.2202, 498.4474,743.61505 |
| TG 46:1 | CID56936638 | C49H92O6 | 20.88 | [M+Na]+ | 799.6785 | 799.6783 | -0.25 | 203.0856,229.1222, 286.2460,524.4674, 580.5285,799.6775 |
| TG 48:2 | CID56936622 | C51H94O6 | 20.87 | [M+Na]+ | 825.6942 | 825.6934 | -0.96 | 229.1213, 522.4606, 579.5186,825.6925 |
| TG 36:0 | CID10851 | C39H74O6 | 18.71 | [M+NH4]+ | 656.5905 | 656.5823 | -12.48 | 229.12682, 257.2108, 411.3468,440.4118,496.4457 |
| TG 57:3;O2 | CID129727532 | C60H110O8 | 19.28 | [M+NH4]+ | 976.862 | 976.8547 | -7.47 | 229.1951, 233.1900, 265.2165, 329.284,705.5988,976.8609 |
| TG 53:3 | CID131754189 | C56H102O6 | 21.84 | [M+NH4]+ | 888.8096 | 888.8012 | -9.45 | 277.2524, 279.2683,617.5499, 871.7741, 888.7931 |
| TG 55:3;O2 | CID166606354 | C58H106O8 | 18.75 | [M+NH4]+ | 948.8307 | 948.8223 | -8.85 | 229.1311, 257.21112:42318 ,311.2565, 678.5744, 948.8214 |
| TG 57:4;O2 | CID9920343 | C60H108O8 | 18.72 | [M+NH4]+ | 974.8464 | 974.838 | -8.61 | 257.2114, 477.3925, 685.5757,703.5864,957.8108, 974.8360 |
| TG 44:1 | CID56936636 | C47H88O6 | 20.31 | [M+Na]+ | 771.6472 | 771.6466 | -0.77 | 203.0855, 211.2058, 229.1202, 552.4968,771.64648 |

[S12].	Kawahata I, Yagishita S, Kazuko H, Nagatsu I, Nagatsu T, Ichinose H. Immunohistochemical analyses of the postmortem human brains from patients with Parkinson’s disease with anti-tyrosine hydroxylase antibodies. 2015.

[S13].	Colamartino M, Padua L, Cornetta T, Testa A, Cozzi R. Recent advances in pharmacological therapy of Parkinson ’ s disease : Levodopa and carbidopa protective effects against DNA oxidative damage. 2012;4:1191–9.

[S14].	Naduthota RM, Bharath RD, Jhunjhunwala K, Yadav R, Saini J, Christopher R, et al. Imaging biomarker correlates with oxidative stress in Parkinson’s disease. Neurol India [Internet]. 2017;65(2). Available from: https://journals.lww.com/neur/fulltext/2017/65020/imaging_biomarker_correlates_with_oxidative_stress.11.aspx

[S15].	Yakunin E, Kisos H, Kulik W, Grigoletto J, Wanders RJA, Sharon R. The regulation of catalase activity by PPAR γ is affected by α-synuclein. Ann Clin Transl Neurol. 2014 Mar;1(3):145–59.

[S16].	Yang W, Chang Z, Que R, Weng G, Deng B, Wang T, et al. Contra-Directional Expression of Plasma Superoxide Dismutase with Lipoprotein Cholesterol and High-Sensitivity C-reactive Protein as Important Markers of Parkinson’s Disease Severity. Front Aging Neurosci. 2020;12:53.

[S17].	Antony PMA, Boyd O, Trefois C, Ammerlaan W, Ostaszewski M, Baumuratov AS, et al. Platelet mitochondrial membrane potential in Parkinson’s disease. Ann Clin Transl Neurol. 2015 Jan;2(1):67–73.

[S18].	Smith AM, Depp C, Ryan BJ, Johnston GI, Alegre-Abarrategui J, Evetts S, et al. Mitochondrial dysfunction and increased glycolysis in prodromal and early Parkinson’s blood cells. Mov Disord. 2018 Oct;33(10):1580–90.

[S19].	Schapira AH, Cooper JM, Dexter D, Jenner P, Clark JB, Marsden CD. Mitochondrial complex I deficiency in Parkinson’s disease. Vol. 1, Lancet (London, England). England; 1989. p. 1269.

[S20].	Arthur CR, Morton SL, Dunham LD, Keeney PM, Bennett JP. Parkinson’s disease brain mitochondria have impaired respirasome assembly, age-related increases in distribution of oxidative damage to mtDNA and no differences in heteroplasmic mtDNA mutation abundance. Mol Neurodegener [Internet]. 2009;4(1):37. Available from: https://doi.org/10.1186/1750-1326-4-37

[S21].	Pérez MJ, Baden P, Deleidi M. Progresses in both basic research and clinical trials of NAD+ in Parkinson’s disease. Mech Ageing Dev [Internet]. 2021;197:111499. Available from: https://www.sciencedirect.com/science/article/pii/S0047637421000713

[S22].	Berven H, Kverneng S, Sheard E, Søgnen M, Af Geijerstam SA, Haugarvoll K, et al. NR-SAFE: a randomized, double-blind safety trial of high dose nicotinamide riboside in Parkinson’s disease. Nat Commun [Internet]. 2023;14(1):7793. Available from: https://doi.org/10.1038/s41467-023-43514-6

[S23].	Doulias PT, Yang H, Andreyev AY, Dolatabadi N, Scott H, K Raspur C, et al. S-Nitrosylation-mediated dysfunction of TCA cycle enzymes in synucleinopathy studied in postmortem human brains and hiPSC-derived neurons. Cell Chem Biol [Internet]. 2023;30(8):965-975.e6. Available from: https://www.sciencedirect.com/science/article/pii/S2451945623001964

[S24].	Willkommen D, Lucio M, Moritz F, Forcisi S, Kanawati B, Smirnov KS, et al. Metabolomic investigations in cerebrospinal fluid of Parkinson’s disease. PLoS One [Internet]. 2018 Dec 10;13(12):e0208752. Available from: https://doi.org/10.1371/journal.pone.0208752
